# Measuring the reproducibility and quality of Hi-C data

**DOI:** 10.1101/188755

**Authors:** Galip Gürkan Yardımcı, Hakan Ozadam, Michael E.G. Sauria, Oana Ursu, Koon-Kiu Yan, Tao Yang, Abhijit Chakraborty, Arya Kaul, Bryan R. Lajoie, Fan Song, Ye Zhang, Ferhat Ay, Mark Gerstein, Anshul Kundaje, Qunhua Li, James Taylor, Feng Yue, Job Dekker, William S. Noble

**Affiliations:** Department of Genome Sciences, University of Washington; Program in Systems Biology, University of Massachusetts Medical School; Biology Department, Computer Science Department, Johns Hopkins University; Department of Genetics, Stanford University; Department of Computational Biology, St. Jude Children’s Research Hospital; Bioinformatics and Genomics Program, Huck Institutes of the Life Sciences, Penn State University; Computational Biology Division, La Jolla Institute for Allergy and Immunology; University of Massachusetts Medical School; Program in Computational Biology and Bioinformatics, Yale University; Department of Computer Science, Stanford University; Department of Statistics, Penn State University; Department of Biochemistry & Molecular Biology, College of Medicine, Penn State University; Howard Hughes Medical Institute

## Abstract

Hi-C is currently the most widely used assay to investigate the 3D organization of the genome and to study its role in gene regulation, DNA replication, and disease. However, Hi-C experiments are costly to perform and involve multiple complex experimental steps; thus, accurate methods for measuring the quality and reproducibility of Hi-C data are essential to determine whether the output should be used further in a study. Using real and simulated data, we profile the performance of several recently proposed methods for assessing reproducibility of population Hi-C data, including HiCRep, GenomeDISCO, HiC-Spector and QuASAR-Rep. By explicitly controlling noise and sparsity through simulations, we demonstrate the deficiencies of performing simple correlation analysis on pairs of matrices, and we show that methods developed specifically for Hi-C data produce better measures of reproducibility. We also show how to use established (e.g., ratio of intra to interchromosomal interactions) and novel (e.g., QuASAR-QC) measures to identify low quality experiments. In this work, we assess reproducibility and quality measures by varying sequencing depth, resolution and noise levels in Hi-C data from 13 cell lines, with two biological replicates each, as well as 176 simulated matrices. Through this extensive validation and benchmarking of Hi-C data, we describe best practices for reproducibility and quality assessment of Hi-C experiments. We make all software publicly available at http://github.com/kundajelab/3DChromatin_ReplicateQC to facilitate adoption in the community.

## Background

The Hi-C assay couples chromosome conformation capture (3C) with next-generation sequencing, making it possible to profile the three-dimensional structure of chromatin in a genome-wide fashion [1]. Recently, application of the Hi-C assay has allowed researchers to profile the 3D genome during important biological processes such as cellular differentiation [2,3], X inactivation [4–6] and cell division [7]; and to identify hallmarks of 3D organization of chromatin, such as compartments [1], topologically associating domains (TADs) [8–10], and DNA loops [11]. Because the Hi-C assay measures the 3D conformation of a genome in the form of pairs of mapped reads (“interactions”) connecting different loci, many such pairs are required to adequately characterize all pairwise interactions across a complete genome [11–13]. Consequently, the Hi-C assay can be costly to run. It is thus essential to have accurate and robust methods to evaluate the quality and reproducibility of Hi-C experiments, both to ensure the validity of scientific conclusions drawn from the data and to indicate when an experiment should be repeated or sequenced more deeply. Reproducibility measures are also important for deciding whether two replicates can be pooled, a strategy that is frequently used to obtain a large number of Hi-C interactions [11].

A rich collection of literature for assessing the quality and reproducibility of a large collection of next generation sequencing based genomics assays, such as ChIP-seq [14] and DNase-seq [15], has been complied over the past decade [16–18]. For these assays, enrichment of signal (“peaks”) at loci of interest [19] and assay-specific properties of sequencing fragments have been used as indicators of the quality of an experiment [16]. Correlation coefficient [20–22] and statistical methods such as the irreproducible discovery rate (IDR) [17] have been used to measure the reproducibility of such assays. However, all of these methods are designed to operate on data that is laid out in one dimension along the genome. Furthermore, unlike other functional genomics assays, Hi-C data must be analyzed at an effective resolution determined by the user [13,23,24]. For these reasons, existing methods for assessing genomic data quality and reproducibility are not directly applicable to Hi-C data.

A variety of methods have been used previously to measure the quality and reproducibility of Hi-C experiments. Ad hoc measures include using, for reproducibility, the Pearson or Spearman correlation coefficient [2,25–27] and, for data quality, statistics that describe the properties of Hi-C fragment pairs [1,28]. The drawbacks of using correlation as a reproducibility measure for genomics experiments, both because of its susceptibility to outliers and because it implicitly treats all elements of the Hi-C matrix as independent measurements, has been documented [16,29]. In practice, because most of the Hi-C signal arises from interactions between loci less than 1 Mb apart [23,24], the correlation coefficient will be dominated by these short range interactions. To alleviate such problems, distance based stratification [30] and dimensionality reduction of Hi-C signal [31], prior to measuring the correlation, have been proposed. Conversely, simple mapping statistics may be used to indicate a high or low percent of invalid or artefactual Hi-C fragments [24,32], but such statistics reflect only the mapping stage of the analysis and cannot be immediately combined into a robust quality score.

To overcome these problems, members of the ENCODE Consortium have recently developed methods for assessing both the quality and the reproducibility of the Hi-C assay [33–36]. In this study, we used large sets of real and simulated Hi-C data to assess and compare the performance of methods for measuring the reproducibility of Hi-C data and evaluating Hi-C data quality. We generated multiple benchmarks for testing the performance of reproducibility measures and established that all of these methods can accurately measure reproducibility of Hi-C data, whereas correlation coefficient cannot. Similarly, we have used real and simulated datasets to profile the performance of quality control methods and compared these methods to established statistics that have been used as indicators of high quality Hi-C experiments. Here, we offer a thorough assessment of quality control and reproducibility methods and describe best practices for analyzing the quality and reproducibility of Hi-C data.

## Results

### Experimental and simulated Hi-C datasets for performance evaluation

We performed two replicate Hi-C experiments on cells from 13 immortalized human cancer cell lines from a variety of tissues and lineages using HindIII and DpnII restriction enyzme digestion (Supp. Table 1). After aligning and filtering of paired end sequencing reads, we obtain 10 to 61 million paired reads per experiment for 11 cell types (generated using HinDIII) and more than 400 millions paired reads for the remaining two deeply sequenced cell types (generated using DpnII). These Hi-C interactions serve as a readout of three dimensional proximity of the corresponding genomic loci. The interactions are binned into fixed-sized bins, and a count of the number of Hi-C interactions that connect each pair of bins is stored in a Hi-C contact matrix. Unless otherwise noted, we used 40 kilobase (kb) bins because this value achieves reasonable sparsity of the Hi-C contact matrices, based on the depth of sequencing of the data sets used in our study. Also, this resolution has been adopted in multiple previous studies [7,8]. We use the resulting Hi-C matrices as input to every reproducibility and quality control analysis in this study, except where indicated.

For use in assessing reproducibility and quality measures for Hi-C data, we designed a model for simulating noisy Hi-C experiments (Fig. 1A). Our noise model aims to simulate a contact matrix from a Hi-C experiment performed on chromatin that lacks any high order structure, such as loops and topologically associating domains. For this purpose, our simulation models two main phenomena: the “genomic distance effect,” i.e., the higher prevalence of crosslinks between genomic loci that are close together along the genome [1], and random ligations generated by the Hi-C protocol [24]. For the first phenomenon, we use real Hi-C data, and we sample from the empirical marginal distribution of counts as a function of genomic distance. The second phenomenon, random ligation noise, is modeled by generating Hi-C interactions between random bin pairs (see Methods for details). Counts generated by these two “noise” components of the model can be mixed with different proportions to produce simulated “pure noise” Hi-C matrices. We then mix the simulated contacts with experimental contact matrices in varying proportions to obtain noise injected matrices.

**Fig 1.**
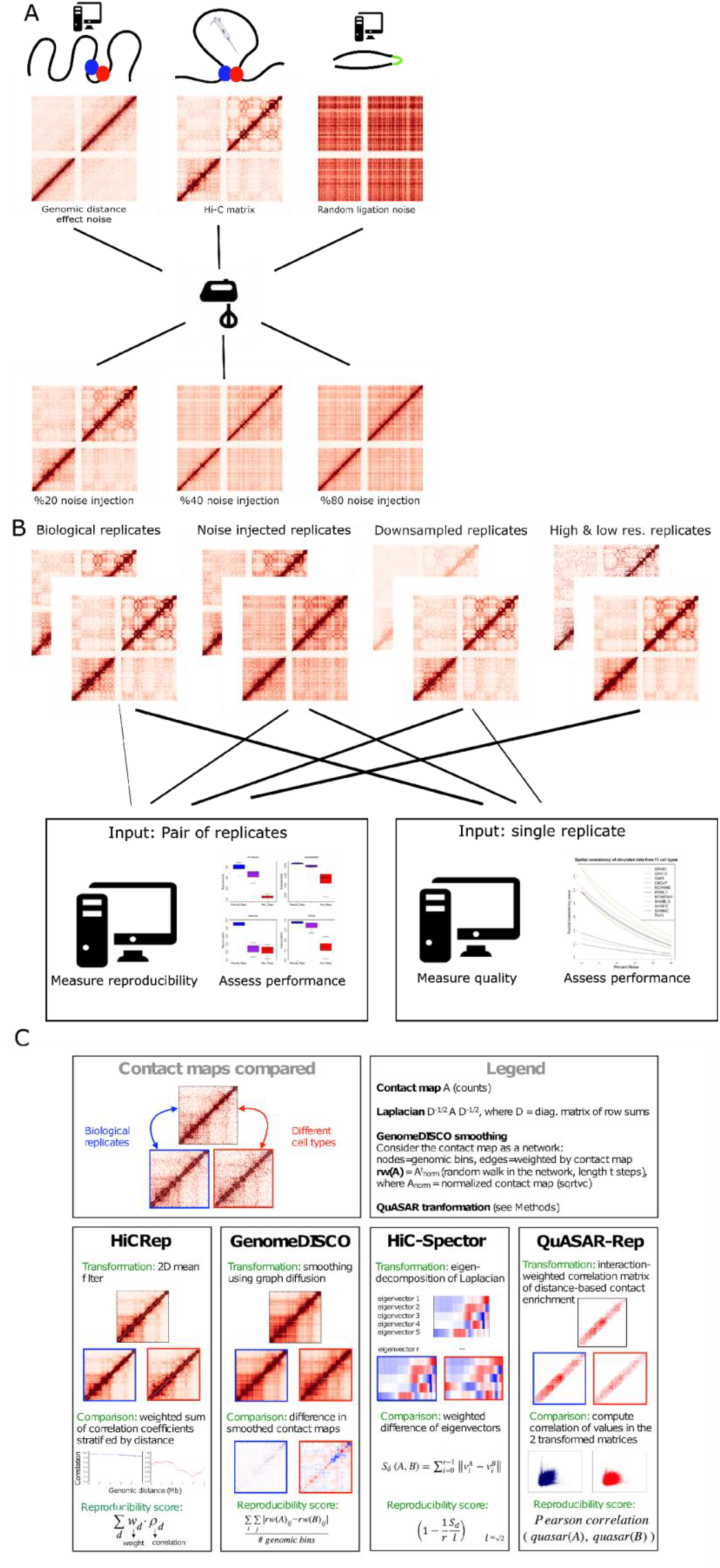
Overview of the study. (A) Schematic showing the approach for generating noise injected Hi-C matrices. In the upper panel, we generate two types of noise from real Hi-C data (center): random ligation noise (right) and genomic distance effect noise (left). The three matrices are then mixed to generate noisy datasets (lower panel). By changing the mixing proportions, we can create datasets with varying percentages of noise. (B) To benchmark the performance of various quality control and reproducibility measures, we compiled a large number of Hi-C replicates from 13 cell types, and simulated noise injected datasets from the original data. Real and simulated datasets binned at different resolutions and downsampled to different coverage levels are the inputs to reproducibility and quality control measures where each replicate pair and single replicate are assigned a score. Performance of each measure is evaluated on their ability to correctly rank real and simulated datasets. (C) Summary of the basic principles of the four reproducibility methods evaluated in this study.

In addition to noise, we tested the effects of sparsity and the resolution of Hi-C matrices on the performance of each method. We profiled the effects of sparsity explicitly by downsampling real Hi-C matrices to contain a set of fixed total number of intra-chromosomal Hi-C interactions. Binning resolution further controls the sparsity of a Hi-C matrix, at the same time dictating the scale of chromatin organization that can be observed in a Hi-C matrix. By binning deeply sequenced Hi-C datasets containing at least 400 million intrachromosomal Hi-C interactions from two cell types, we generated Hi-C matrices binned at high, mid and low resolutions (10 kb, 40 kb, 500 kb) and used these to investigate the effect of resolution on each method as well (Supp. Table 1). A schematic of the full range of datasets used in this study to validate each method is shown in Fig.1B.

### Measures for quality and reproducibility of Hi-C data

Four recently developed methods for measuring the quality of and reproducibility of Hi-C experiments were assessed in this study (Fig. 1C). HiCRep [34], GenomeDISCO [35], HiC-Spector [33] and QuASAR-Rep [36] measure reproducibility, and QuASAR-QC measures quality of Hi-C data. The four reproducibility methods we evaluate employ a variety of transformations of the Hi-C contact matrix. HiCRep stratifies a smoothed Hi-C contact matrix according to genomic distance and then measures the weighted similarity of two Hi-C contact matrices at each stratum. In this way, HiCRep explicitly corrects for the genomic distance effect and addresses the sparsity of contact matrices through stratification and smoothing, respectively. GenomeDISCO uses random walks on the network defined by the Hi-C contact map to perform data smoothing before computing similarity. The resulting score is sensitive to both differences in 3D DNA structure and differences in the genomic distance effect [35], and makes it thus more challenging for two contact maps to be reproducible, as they have to satisfy both criteria to be deemed similar. HiC-Spector transforms the Hi-C contact map to a Laplacian matrix and then summarizes the Laplacian by matrix decomposition. QuASAR calculates the interaction correlation matrix, weighted by interaction enrichment. The two variants of QuASAR, QuASAR-QC and QuASAR-Rep, both assume that spatially close regions of the genome will establish similar contacts across the genome, and they measure quality and reproducibility, respectively, by testing the validity of this assumption for a single and pair of replicates.

### Reproducibility measures correctly rank noise injected datasets

To assess the performance of the reproducibility measures, we simulated pairs of Hi-C matrices with varying noise levels. Intuitively, a good reproducibility measure should declare the least noisy replicate pair as most reproducible and the noisiest replicate pair as least reproducible. We paired a real Hi-C contact matrix with a noisier version of the same matrix using a wide range of simulated noise levels (5%, 10%, 15%, 20%, 30%, 40% and 50%). This procedure yielded seven pairs of replicates for each of 11 different cell types. We performed this approach using two different sets of randomly generated noise matrices, using one-third genomic distance noise and two-thirds random ligation noise or vice versa. Each replicate pair was assigned a reproducibility measure by HiCRep, GenomeDISCO, HiC-Spector, QuASAR-Rep and Pearson correlation.

Our analysis showed that all reproducibility measures were able to correctly rank the simulated datasets. Averaged over 11 different cell types, we observed a monotonic trend for all of these measures (Fig. 2A). Indeed, for every cell type and every measure, increasing the noise level always led to a decrease in estimated reproducibility (Supp. Fig. 1). Qualitatively, the trends in Figure 2A suggest that QuASAR and HiCRep may be more robust to noise than the other reproducibility measures.

**Figure 2.**
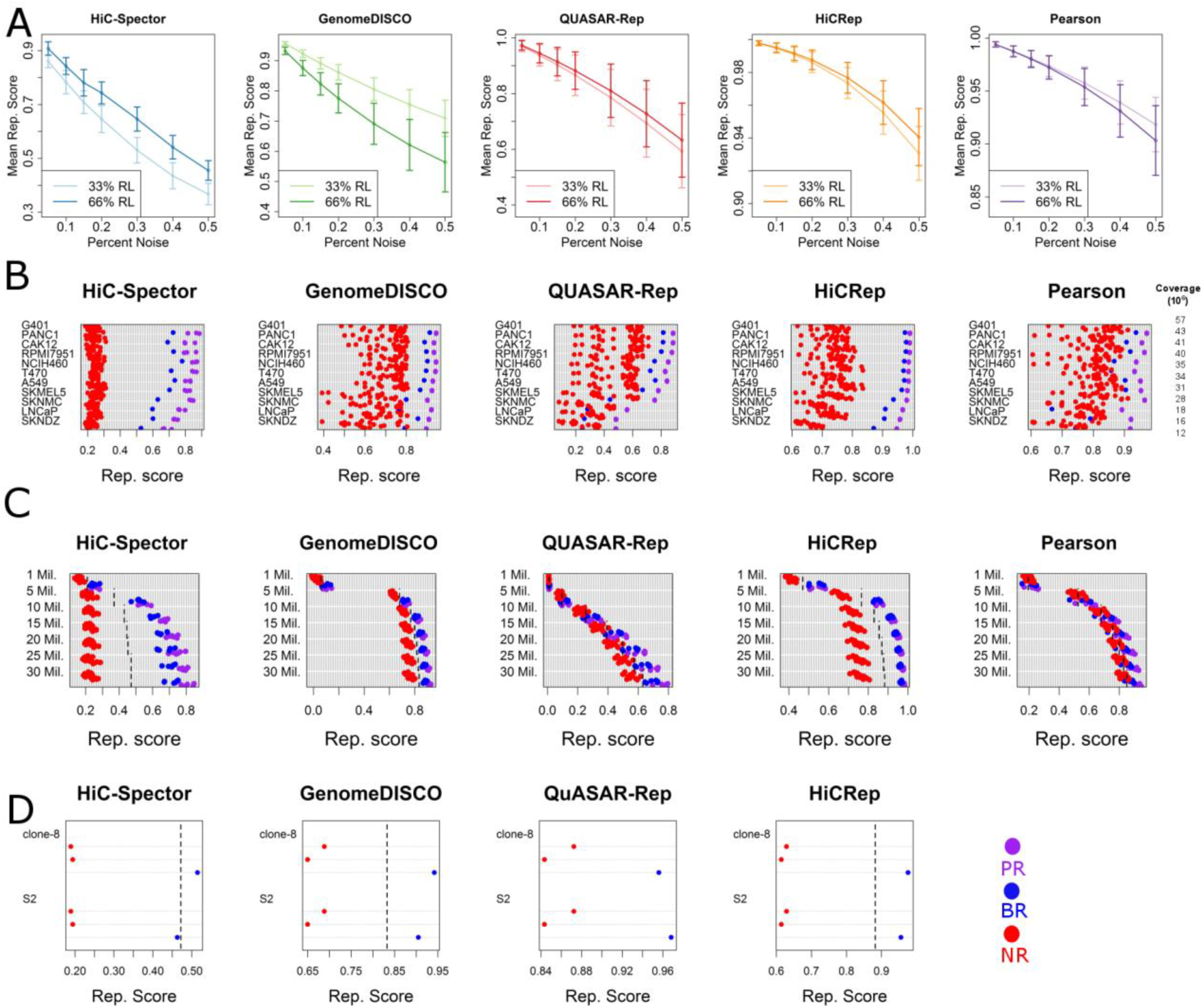
Comparison of reproducibility measures. (A) Curves showing the mean reproducibility score assigned to 11 cell types at each noise injection level for 33% and 66% random ligation noise configurations. Vertical bars represent one standard deviation away from the mean. (B) Reproducibility scores assigned to biological replicate (blue), non-replicate (red) and pseudo-replicate (purple) pairs for each cell type. Coverage values are the mean number of interactions for each pair of replicates. (C) Reproducibility scores assigned to biological replicate (blue), non-replicate (red), and pseudo-replicate (purple) pairs from six cell types at seven different coverage levels. Dashed lines indicate the empirical threshold for distinguishing biological replicate pairs from non-replicate pairs. (D) Reproducibility scores assigned to biological replicate (blue) and non-replicate (red) pairs for clone-8 and S2 cells from *Drosophila*. Each panel shows the separation between two replicate pair types for each Hi-C reproducibility measure. Dashed lines correspond to the empirical thresholds inferred from human Hi-C data.

Comparing the two noise models, we saw less consistent trends. HiC-Spector assigned higher reproducibility scores to matrices with 66% genomic distance noise and 33% random ligation noise. GenomeDISCO showed the opposite behavior whereas QuASAR-Rep, HiCRep and Pearson correlation gave similar scores regardless of the underlying noise proportions. This variability suggests that the various reproducibility measures exhibit different sensitivities to different sources of noise, thus potentially yielding complementary assessments of reproducibility.

### Assessment using real data sets reveals differences among reproducibility measures

Inevitably, any simulation approach is only as good as its underlying assumptions; thus, we also analyzed the performance of the four reproducibility measures using real data. Specifically, we asked whether the reproducibility measures can discriminate between pairs of independent Hi-C experiments repeated on the same cell type versus pairs of experiments from different cell types. In this setup, we used three types of replicate pairs: a single pair of matrices from the same cell type (which we call “biological replicates,” although each pair represents the same cells being prepped twice, rather than two different sets of cells), pairs of matrices from different cell types (non-replicates) and pairs of matrices sampled from combined biological replicates (pseudo-replicates, see Methods for details about the generation of pseudo-replicates) [34]. We assigned a reproducibility score to every matrix pair for each measure and asked if reproducibility scores differ among replicate pair types.

Because pseudo-replicates are generated from pooled biological replicates, their variation solely stems from statistical sampling, with no biological (including distance effect) or technical variance. Therefore, we expect pseudo-replicates to exhibit the highest reproducibility. Conversely, non-replicate pairs are expected to have the lowest degree of reproducibility, because they contain all the experimental variation observed in biological replicates, as well as cell type specific differences in 3D chromatin organization.

In contrast to the simulation analysis, the analysis using real datasets showed distinct differences among the five methods. For each of the eleven cell types and each reproducibility measure, we assigned reproducibility scores to a single biological replicate pair, 20 non-replicate pairs, and three pseudo-replicate pairs (Fig. 2B). The reproducibility score of a replicate pair is the score obtained by averaging reproducibility scores assigned to each chromosome. All four reproducibility measures and the Pearson correlation can separate replicate pair types from each other (Supp. Fig. 2); however, the reproducibility measures generally achieved clearer separation between different replicate pair types. These differences are statistically significant according to a one-sided Kolmogorov-Smirnov test (P < 0.01). In addition to the Pearson correlation, we considered the rank-based Spearman correlation as a potential method for assessing reproducibility. We also considered using either type of correlation in conjunction with ICE normalization. The results (Supp. Fig. 3) show that none of these four methods successfully separates biological replicate from non-replicate pairs. Intuitively, we prefer a measure that separates non-replicates from biological replicates with a clear margin. By this measure, the Hi-C-Spector measure yields the largest separation, followed by HiCRep, QuASAR-rep and GenomeDISCO (Fig. 2B). Among them, HiC-Spector and HiCRep correctly rank all replicates types for all eleven comparisons, with a clear separation between biological replicates and non-replicates. GenomeDISCO ranks a biological replicate lower than a non-replicate for a single case out of eleven. The pair of biological replicates that GenomeDISCO ranks lower than non-replicates shows a marked difference in genomic distance effect (Supp. Fig. 4), to which this method is sensitive [35]. QuASAR-rep is able to correctly rank biological replicates above non-replicates in seven out of eleven cases. The cell types in which it fails have only 12 to 28 million interactions, suggesting that QuASAR-Rep does not perform well when coverage is low and the resolution is set to 40 kb. However, reanalysis of the same data suggests that switching to a larger resolution (120 kb) improves QuASAR-rep’s performance, leading to separation between replicates and non-replicates for all cell lines but two (data not shown). As expected, the Pearson correlation performs worse than the Hi-C-specific measures, ranking non-replicates higher than biological replicates in seven cases.

Pseudo-replicate reproducibility scores provide an upper bound for each reproducibility measure. In general, these scores show similar trends to those described above. For example, the Pearson correlation scores assigned to pseudo-replicates show a relatively wide separation from the rest of the scores, even though non-replicates and biological replicates are intermingled. On the other hand, GenomeDISCO, HiC-Spector, HiCRep, and QuASAR-rep show the desired behaviour: a high degree of separation between non-replicates and biological replicates, and a relatively small separation between biological replicates and pseudo-replicates.

### Reproducibility can be determined over a range of experimental coverage

To directly investigate the effects of the coverage of a Hi-C experiment on the reproducibility measures, we downsampled real Hi-C matrices to contain fewer interactions and examined the effects on the resulting reproducibility scores. We limited this analysis to real data from six cell types with higher coverage, and we subsampled each replicate multiple times to contain 1 to 30 million total Hi-C interactions (see Methods for details). These datasets were used for testing the ability of each method to distinguish among different replicate types at lower coverage levels, and for explicitly profiling the dependence of reproducibility scores on coverage levels.

Hi-C reproducibility measures retained their ability to distinguish between replicate types, even at extremely low coverage levels. Visualization of the reproducibility scores revealed that the HiCRep, HiC-Spector and GenomeDISCO measures successfully separate non-replicates from biological replicates even with only five million Hi-C interactions, a feat that Pearson correlation cannot achieve at even the highest coverage level (Fig. 2C). QuASAR-rep can successfully separate biological replicates from non-replicates at 25 and 30 million interactions but fails to distinguish them when coverage is lower than 20 million interactions, consistent with the results from Figure 2B. As before, pseudo-replicate pairs continue to serve as an upper bound for reproducibility measures. However, the separation between pseudo-replicates and biological replicates is reduced at lower coverage levels, and so is the separation between biological replicates and non-replicates. Furthermore, this analysis suggests we can infer empirical thresholds for these reproducibility measures that can effectively separate all biological replicates from non-replicates at a given coverage level, as explained in methods section. These empirical thresholds, selected as the midpoint between the most reproducible non-replicate pair and the least reproducible replicate pair, are shown as dashed lines in Fig. 2C and can be found in Supplemental Table 2.

Consistent with the trends observed in the analysis of real datasets, the reproducibility of downsampled replicate pairs exhibits a dependence on sequencing depth. We observe that reproducibility scores associated with biological replicates become significantly smaller as coverage decreases, according to a one-sided Wilcoxon signed rank test (P < 0.05, Supp. Fig. 5). The HiCRep, GenomeDISCO, QuASAR-rep and Pearson correlation scores exhibit a statistically significant drop for every level of coverage. In contrast, reproducibility scores from HiC-Spector only start to significantly and consistently decay below 20 ×10^6^ interactions, exhibiting a lesser degree of dependence on the coverage level. This may be because the leading eigenvectors used by HiC-Spector tend to capture local or mesoscopic structures, which are less likely to be affected by coverage. Despite varying levels of dependence on coverage, downsampling analysis convincingly shows that all measures exhibit a dependence on coverage. Thus, coverage of different replicate pairs must be factored into reproducibility analyses, especially for comparative purposes.

### Reproducibility measures are robust to changes in resolution

The resolution of a Hi-C matrix effectively dictates the scale of 3D organization observable from the data: a low resolution matrix can only reveal compartments and TADs [1,8], whereas high resolution matrices reveal additional finer scale structures like chromatin loops [11]. To investigate the effect of resolution on reproducibility, we used deeply sequenced Hi-C replicates with at least 400 million intra-chromosomal interactions generated from the HepG2 and HeLa cell lines. From these data, we generated real and simulated replicate pairs at 10kb, 40kb and 500kb resolution, and we measured the reproducibility of each replicate pair.

HiCRep, GenomeDISCO, HiC-Spector, QuASAR-Rep and Pearson correlation accurately measure reproducibility at both high and low resolutions. The four Hi-C-specific methods can correctly rank pseudo, biological and non-replicate pairs at 10kb, 40kb and 500kb resolutions (Fig. 3A) with a clear margin between biological replicate and non-replicate pairs. Surprisingly, we found that the Pearson correlation can correctly rank replicate types for these deeply sequenced datasets. Notably, the reproducibility scores from the four methods are largely independent of resolution. While GenomeDISCO and especially QuASAR-rep exhibit some dependence of resolution, assigning lower reproducibility scores to replicates with lower coverage, they maintain a clear boundary with large margins between biological and non-replicates at all resolutions. However, the Pearson correlation exhibits a larger degree of dependence on resolution for all replicate pair types and maintains relatively smaller margins between non-replicate and biological replicate pairs. Simulated datasets further validate that reproducibility scores from each method decrease with increasing levels of noise at 10kb, 40kb and 500kb resolution (Fig. 3B)

**Fig 3.**
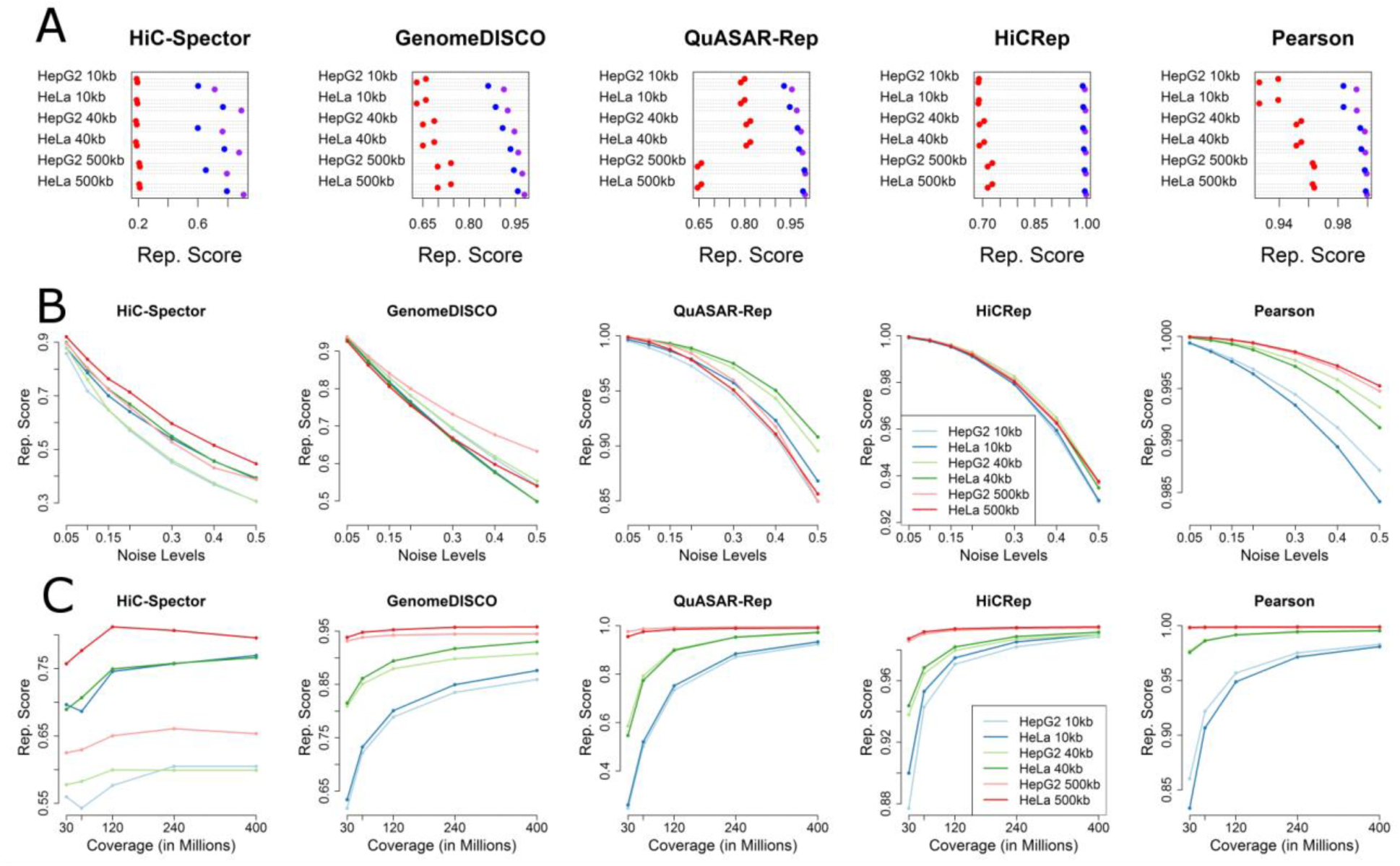
Effects of resolution on reproducibility measures. (A) Reproducibility scores assigned to biological replicate (blue), non-replicate (red), and pseudo-replicate (purple) pairs from HepG2 and HeLa Hi-C datasets at 10 kb, 40 kb and 500 kb resolutions. (B) Reproducibility scores assigned to different cell types at different resolutions, plotted as a function of noise level. (C) Reproducibility scores assigned to downsampled biological replicate pairs at different resolutions. Both the HepG2 and HeLa data sets contain >400 million read pairs.

Next, we used deeply sequenced datasets to further investigate the effect of coverage on reproducibility scores of biological replicates at three resolution levels using a wider range of coverage values (30, 60, 120, 240, and 400 million intra-chromosomal interactions). For HiCRep, QuASAR-rep and GenomeDISCO, we observed that reproducibility scores tend to plateau at 240 million interactions at 10kb and 40kb resolutions, whereas reproducibility scores of 500kb resolution matrices benefit little from higher coverage (Fig. 3C). Consistent with our previous observations, HiC-Spector exhibits a lower degree of dependence on coverage, with scores reaching maxima at 120kb. Overall, the four Hi-C reproducibility measures exhibit robustness to coverage and resolution differences, as measured by their ability to distinguish between replicate and non-replicate pairs.

Next we tested whether the reproducibility measures can be used to select empirically the optimal resolution for a Hi-C dataset. Although resolution strongly influences almost every downstream analysis of Hi-C data, this parameter is generally set in an *ad hoc* fashion. To explore the performance of the measures as a function of the resolution parameter, we binned four pairs of biological replicates at increasingly high resolution ranging from 40 kb, 20 kb, 10 kb and 5 kb and asked if the reproducibility scores of biological replicates decay significantly at higher resolutions. We chose six samples performed using HindIII with coverage values ranging from 15 million to 60 million interactions and two samples generated using DpnII and coverage of ~400 million interactions.

We observed that the four reproducibility measures show variable trends in how reproducibility scores assigned to biological replicates decay with respect to increasing resolution (Supp. Fig. 6). For HiCRep, GenomeDISCO and QuASAR-Rep, the HindIII replicates (A549, G410 and LNCaP) exhibit a decay in reproducibility scores, whereas the scores assigned to replicate pairs generated by DpnII (HepG2) are more robust to changes in resolution. Notably, for these three reproducibility measures, the degree of decay also correlates with the sequence coverage of the data. For HiC-Spector, we do not observe consistent trends. These observations generally support the idea that deeply sequenced replicates generated by a 4-cutter such as DpnII can support resolutions higher than 40 kb, whereas relatively shallow replicates (<100 million read pairs) generated using a 6-cutter are not suitable for binning resolutions higher than 40kb. However, given the lack of a clear elbow or maximum in Supplemental Figure 6, we do not recommend using reproducibility scores to attempt to select an appropriate resolution.

Finally, we compared the run times of each reproducibility measure, using a large number of pairs of chromosome 21 contact matrices binned at 40 kb resolution. As seen in Supplemental Figure 7, QuASAR-Rep achieves the fastest median running time (0.82 s), followed by HiC-Spector (2.76 s), GenomeDISCO (5.77 s) and HiCRep (9.00 s).

### Reproducibility measures accurately quantify reproducibility of Hi-C data from non-human genomes

We investigated whether the four Hi-C reproducibility measures can be applied to data derived from a non-human genome. We wanted to investigate a genome that is markedly different from human, but replicate Hi-C experiments in organisms other than human and mice are rare. We used Hi-C data from Ramirez et al, which has two biological replicates from two cell types (clone-8 and S2) from the fruitfly *Drosophila melanogaster* [37]. The fruitfly genome is approximately 18 times smaller than the human genome. For this analysis, we binned the Hi-C matrices at 10 kb and compared the reproducibility of the four large, non-heterochromatic chromosomes in *Drosophila* (chromosomes 2, 3, 4 and X). As before, we assigned reproducibility scores to each replicate pair and each non-replicate pair. The results show that biological replicate pairs are clearly separated from non-replicates for each measure in both cell types (Figure 2D). Furthermore, for three out of the four reproducibility measure, the empirical thresholds that we inferred from the human Hi-C data (shown as dashed lines in Fig 2D) generalize to the fruitfly genome.

### Noise reduces the consistency and the prevalence of higher order structures in Hi-C matrices

Having investigated four different methods for evaluating the reproducibility of a given pair of Hi-C matrices, we now focus on methods for evaluating the quality of a single HiC matrix. As before, we perform this evaluation by injecting noise into real Hi-C data, producing a collection of 88 matrices corresponding to 11 cell types and 8 different noise profiles (see Methods). Among our four Hi-C reproducibility measures, only one (QuASAR-QC) provides a variant to assess the quality of a single matrix. The procedure yields a single, bounded summary statistic indicative of homogeneity of the underlying sample population and the signal-to-noise ratio of the interaction map. In addition to QuASAR-QC analysis, we profiled two well-known features of 3D organization: statistically significant long range contacts [38,39], which include DNA loops, and topologically associating domains (TADs). Intuitively, we expect that significant contacts and TADs should be harder to detect in noisy matrices, and that such matrices should have a lower degree of consistency.

Our analysis suggests that QuASAR-QC is indeed sensitive to the noise and the coverage of a Hi-C matrix. For each simulated Hi-C matrix from 11 cell types, QuASAR-QC detects a perfectly monotonic relationship between the noise level and the consistency of the matrix (Fig. 4A). The same trend is observed in deeply sequenced HepG2 and HeLa cell types at 10kb, 40kb and 500kb resolutions (Supp. Fig. 8). Although the majority of noise-free combined replicates are assigned a QuASAR-QC score ranging from 0.05 to 0.07, three cell types have strinkingly lower QuASAR-QC scores ranging from 0.03 and 0.02. The Hi-C matrices from these three cell types (LNCaP, SKNDZ, SKNMC) contain fewer Hi-C interactions. Thus, the lower consistency scores are likely partially due to the sparsity that results from low experimental coverage (Supp. Table. 1). Furthermore, investigation of contact probabilities at given genomic distances for each cell type revealed that the three cell types with lower QuASAR-QC scores have significantly higher contact probabilities at genomic distances larger than 50 megabases (Supp. Fig. 9). Because such long range contacts are unlikely to occur due to the organization of chromatin, it is likely that such long range contacts represent random ligation of uncrosslinked DNA fragments, which is a known source of noise in a Hi-C experiment [24]. Thus, the QuASAR-QC measure is potentially sensitive to both the level of simulated noise and the differences in level of inherent noise that each combined replicate contains.

**Figure 4.**
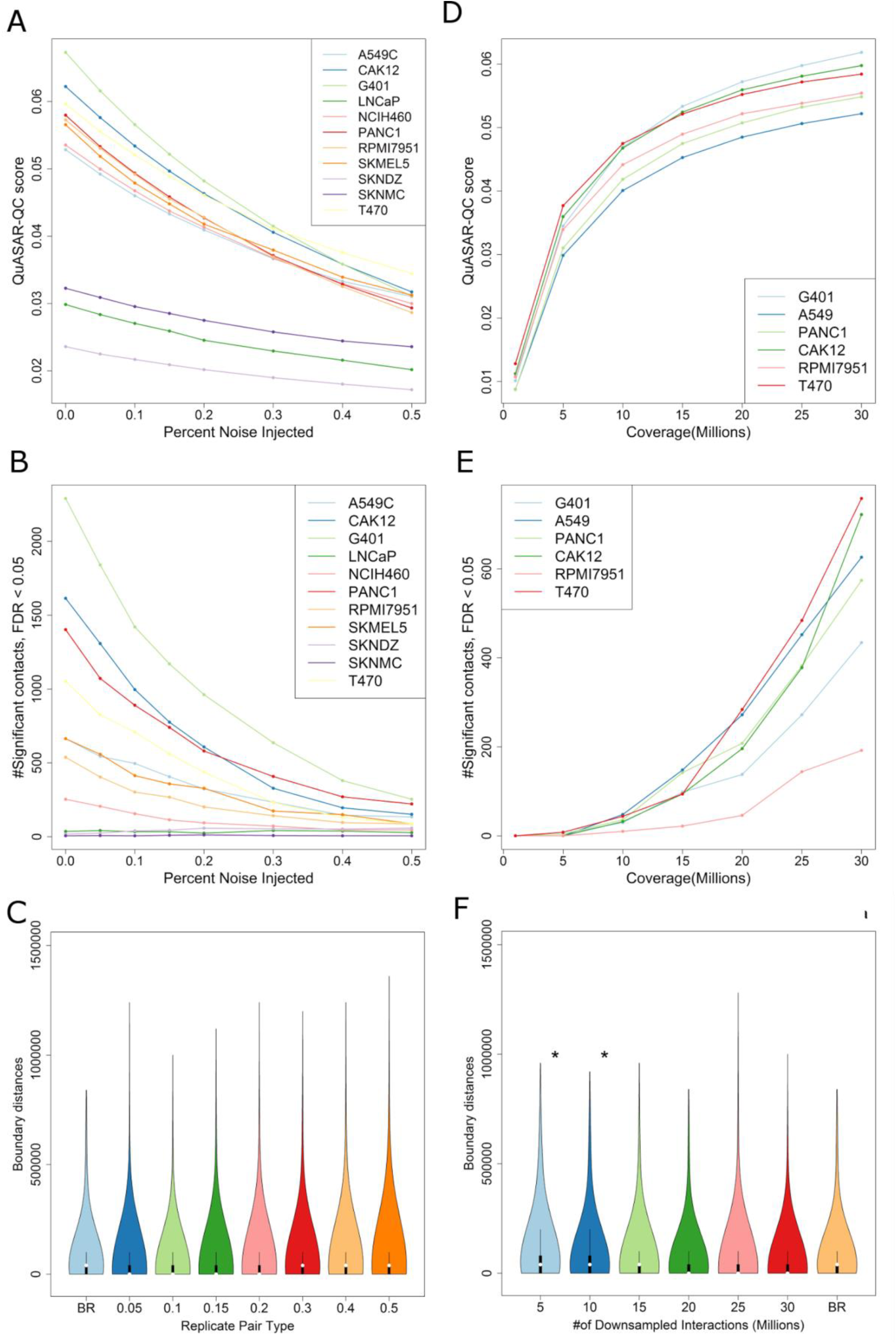
Quality measures. (A) QuASAR-QC scores assigned to noise injected matrices from 11 cell types (B). Total number of significant contacts above a 5% FDR threshold from noise injected matrices from 11 cell types. (C) Violin plots showing the distribution of TAD boundary distances between biological replicates and noise injected replicates for T470 cells. There is no significant change in the distribution of TAD boundary distances at any given noise level. (D) QuASAR-QC scores assigned to downsampled replicates from six different cell types. (E) Total number of significant contacts above a 5% FDR threshold from downsampled replicates from six different cell types. (F) Violin plots showing the distribution of distances between domain boundaries in biological replicates and noise injected replicates for T470 cells. In panels C and F, asterisks indicate that the distribution of boundary distances is significantly larger than the null distribution, which is obtained by comparing biological replicates.

Statistically significant mid-range (50kb-10Mb) interactions are depleted in noisy Hi-C matrices. We identified statistically significant Hi-C contacts using Fit-Hi-C [38] for each of the Hi-C matrices that make up our simulated dataset. Because robust identification of such contacts requires deeply sequenced datasets that contain large numbers of Hi-C interactions, we chose to use a somewhat liberal false discovery rate threshold of 0.05 to facilitate discovery of statistically significant contacts. For 11 cell types, we observed that eight out of eleven cell types exhibit a perfect or near perfect anticorrelation between the injected noise percentage and the total number of significant interactions (Fig. 4B). For the other three cell lines (LNCaP, SKNDZ, SKNMC), Fit-Hi-C identifies almost no significant contacts with or without any noise injection, further supporting the conclusion that these Hi-C data sets have low quality. These three cell lines are also the cell lines that have the lowest QuASAR-QC scores, corroborating the results between these two independent analyses. For the deeply sequenced two data sets (HepG2 and HeLa), we observed a similar trend at both 10kb and 40kb resolutions, with a higher number of significant mid-range contacts due to the higher coverage, as expected (Supp. Fig. 10).

Surprisingly, we found that topologically associating domain detection is highly robust to noise. We identified TADs using the insulation score [5,40] method for the 88 simulated matrices, and we characterized the changes in total number of TADs and TAD size distribution and the changes to TAD boundaries with respect to noise level. The total number of identified TADs and their size distribution are only altered at the highest level of noise injection (Supp. Fig. 11 and 12). In addition, TAD boundaries between the original replicate and noise injected levels exhibit the same degree of variation between two biological replicates, further supporting the idea that TAD boundaries identified with the insulation score approach are highly robust to noise (Fig. 4C, Supp. Fig. 13).

### Quality control measures require different levels of experimental coverage

Continuing our assessment of Hi-C quality measures, we used downsampled Hi-C matrices to investigate the relationship between experimental coverage and each QC measure using a similar setup as before (see Methods).

Quality control metrics exhibit a predictable dependence on the coverage of Hi-C matrices. For each of the six cell types we downsampled, we observed that QuASAR-QC scores are lower for Hi-C matrices with fewer interactions (Fig. 4D). We observe the same trend for deeply sequenced matrices at 10kb and 40kb resolutions; however, QuASAR-QC scores at 500kb tend to benefit less from deeper coverage, likely because coarse resolutions do not require large numbers of Hi-C interactions. (Supp. Fig. 14). Similarly, the number of statistically significant long range interactions also decreases as we reduce the number of total Hi-C interactions. However, the number of significant interactions decrease at a much higher rate: even at 15 million interactions most cell lines lose the majority of significant interactions (Fig. 4E). Larger numbers of significant interactions are detected in deeply sequenced datasets, due to added statistical power, but a similar relationship between coverage and number of significant contacts is observed at both 10kb and 40kb resolutions (Supp. Fig. 15). Conversely, we found that TADs detected by insulation score are robust to low coverage levels. Using the same approach for noise injected datasets, we found that total number of TADs and their size distribution are not altered by lower coverage (Supp. Fig. 16 and 17). Indeed, the distances between TAD boundaries identified at lower coverage and original replicates only differ from the baseline distribution at 10 million or fewer interactions (Fig. 4F, Supp. Fig. 18).

### Quality control measures are consistent with mapping statistics

To further validate the performance of the quality controls measures at our disposal, we investigated the relationship between the QuASAR-QC scores assigned to real Hi-C matrices and various read-mapping statistics that have been used previously to evaluate Hi-C data quality [24]. The four statistics we compared against are the percentages of fragment pairs that can be mapped uniquely to the genome (aligned pairs), fragment pairs from the same restriction fragments (invalid pairs), intrachromosomal interactions (intra-chromosomal percentage), and fragment pairs that are repeated in the data set (PCR duplicate rate).

Overall, we observe varying degrees of correlation between the quality control measures and the mapping statistics for biological replicates. The percentage of aligned pairs is correlated with higher quality experiments, consistent with what one would intuitively expect from high quality sequencing libraries (Fig. 5A). The percentage of invalid pairs is also weakly anti-correlated with QuASAR-QC scores, consistent with the fact that invalid pairs represent uninformative Hi-C interactions (Fig. 5B). However, we observed the highest degree of correlation between QuASAR-QC scores and intrachromosomal percentage (Fig. 5C). In a typical Hi-C experiment, a portion of inter-chromosomal interactions result from random ligation of non-crosslinked fragments; thus, a significant enrichment of inter-chromosomal interactions, which results in a depletion of intra chromosomal interactions, indicates a low quality Hi-C experiment. In particular, six biological replicates with lower than 30% intra-chromosomal interactions have the lowest QuASAR-QC scores; these replicates are from the LNCaP, SKNDZ, and SKNMC cell types. Analysis of downsampled data shows that this effect is not simply due to the overall lower sequencing depth of these three replicates (Supp. Fig. 19). These replicates were also identified to have lower quality in our simulation studies (Fig. 4A) and are depleted for significant mid-range interactions, establishing the consistency of quality control measures overall. We note that this finding is consistent with the previously suggested range of 40-60% intra-chromosomal interactions for high quality experiments [24]. The PCR duplicate rate is uncorrelated with QuASAR-QC.

**Figure 5.**
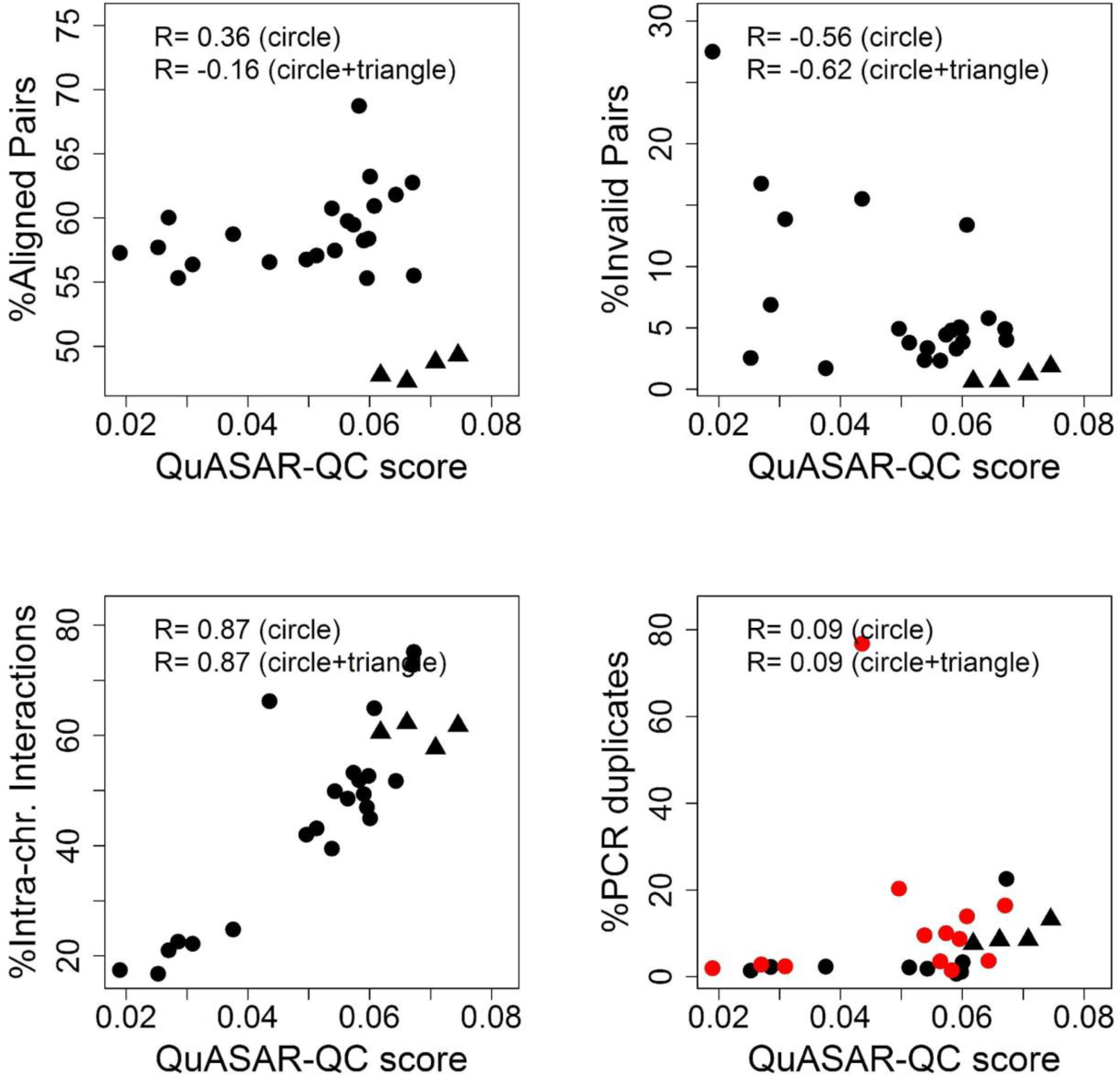
Comparison of QuASAR-QC to mapping statistics. Scatter plots of QuASAR-QC scores of biological replicates from 13 cell types plotted against quality statistics that describe percentages of (A) successful mapping, (B) artifactual Hi-C fragments, (C) intra-chromosomal interactions, and (D) PCR duplicates. Dots correspond to low coverage Hi-C replicates from 11 cell types generated using HindIII, and triangles correspond to replicates from two deeply sequenced cell types generated by DpnII. Red dots correspond to a subset of samples with very similar total coverage (138-171 million read pairs). Each plot lists two Pearson correlation coefficients: the correlations between the given statistic and QuASAR-QC scores for only the 11 HinDIII cell types and for all 13 cell types.

Note that the PCR duplicate rate may be influenced by overall coverage, which we have not controlled for in this experiment. Nonetheless, even for sets of experiments with very similar coverage (red dots in Figure 5D), we observe very little correlation

## Discussion

We evaluated recently proposed methods for measuring the quality and reproducibility of Hi-C experiments. Using a rich set of Hi-C experiments from a variety of human cell types, we tested whether these methods can identify reproducible and high quality experiments. Furthermore, we generated Hi-C contact matrices with controlled levels of noise by designing a simulated noise injection process. Our analysis shows that these measures perform well and improve upon the shortcomings of using generic or qualitative approaches.

The Hi-C reproducibility measures that we evaluated assess reproducibility more accurately than the Pearson or Spearman correlation for real and simulated datasets. In particular, measures specifically designed for Hi-C data can better distinguish subtle differences in the 3D organization of different cell types, because these methods directly account for the special noise properties of this data type that are overlooked by traditional similarity scores.

Selecting an appropriate reproducibility measure for a given study may depend in part upon the goals of the study. A scientist may be primarily interested in a measure that distinguishes between biological replicates and non-replicates. Such a goal might be appropriate, for example, if the method will be used to check for sample swaps during large-scale experiments. In this setting, our results show that HiC-Spector often had the best margin among all four measures (Figure 2B and C). This is true even when we place all four measures on a similar scale by using the variance associated with non-replicate pairs (data not shown). On the other hand, simply discriminating among replicates and non-replicates may not be sufficient in some contexts. If the study aims to use the reproducibility measure to quantify similarities among various experiments, then HiCRep has been shown previously to discriminate well among cell types [34], whereas the other methods in this study have not been examined in this fashion.

Furthermore, our analysis also suggests that the different reproducibility measures may be more sensitive to different types of noise, with GenomeDISCO showing more sensitivity to random ligation noise than to genomic distance noise, HiC-Spector showing the opposite behavior, and QuASAR-Rep and HiCRep showing similar sensitivities to both types of noise (Figure 2A). Because genomic distance noise preferentially affects short range Hi-C interactions, this observation is consistent with the hypothesis that HiC-Spector largely focuses on local structures which are detected by short range Hi-C interactions. Overall, QuASAR-Rep and HiCRep appear to exhibit an overall lower sensitivity to varying noise levels than HiC-Spector and GenomeDISCO. Also, GenomeDISCO tends to be more sensitive to differences in genomic distance effect between the samples compared [35].

The scores produced by all four reproducibility methods decrease in the presence of decreasing sequencing depth and fixed resolution or in the presence of increasing resolution at a fixed sequencing depth. Nonetheless, three out of four methods (GenomeDISCO, HiCRep, and HiC-Spector) show robustness to increasing sparsity, as measured by their ability to distinguish replicate from non-replicate pairs. Only QuASAR-Rep fails to measure reproducibility accurately for the most sparse datasets at high resolutions, though this effect is ameliorated if the data is analyzed using a lower resolution (Data not shown). Thus, we hypothesize that one reason why GenomeDISCO and HiCRep perform well on low coverage data sets is because they perform smoothing on the contact matrix. Overall, these results suggest that experimenters can assess whether a given set of samples are “reproducible enough” with as few as valid five million Hi-C interactions and then follow up with deeper sequencing. Among the four methods, HiC-Spector exhibits the least dependence on sequencing depth (Figure 3C) or resolution (Supp. Fig. 6). These results are further consistent with the hypothesis that HiC-Spector focuses on local features of chromatin structure, which explains HiC-Spector’s robustness to low coverage.

Note that if the goal of a study is to quantify similarities among various experiments, then the dependence of reproducibility scores on data sparsity must be taken into account. For example, in our study, the SKMEL5 and SKNMC experiments differ in sequencing depth by a factor of 2. This difference could confound attempts to cluster or hierarchically organize cell types. In such a setting, all data sets should be randomly downsampled to a common sequencing depth prior to analysis.

An important question is whether the methods and thresholds derived here will generalize to non-human genomes. Preliminary analysis (Figure 2D) suggests that the empirical reproducibility thresholds derived for GenomeDISCO, HiCRep and HiC-Spector may generalize to the much smaller Drosophila genome, whereas the QuASAR-Rep measure does not. However, this result is preliminary due to the small number of currently available, replicated Hi-C experiments in non-human genomes.

The QuASAR-QC measure provides an interpretable score that can accurately rank simulated datasets according to noise levels and distinguish low quality real Hi-C experiments from high quality ones (in submission). This measure correlates with previously established statistics that indicate high quality in a Hi-C experiment and have been used as qualitative indicators of quality. Each of these statistics captures different sources of error in a Hi-C assay. In contrast, QuASAR-QC offers a single score that allows direct ranking of multiple experiments.

Significant mid-range interactions, such as DNA loops, are also depleted in low quality Hi-C experiments in both simulated and real datasets. Surprisingly, we found that TAD detection is fairly robust to all but high levels of noise, presumably because TAD detection only requires that a dataset contains a sufficient proportion of valid short range Hi-C interactions and ignores mid‐ and long-range interactions. Unfortunately, it is challenging to convert the enrichment of such features into a quality control measure, due to other quality-independent biological processes which can cause variation of these features. However, a near total depletion of these features, mid-range interactions in particular, may certainly indicate lower quality overall.

We anticipate that the reproducibility measures we evaluated in this study may be applicable to data from recently developed single cell Hi-C assays [41–43]. The primary challenge, in this setting, would be the extreme sparsity of single cell data. Our experiments show that, even when we randomly downsample to 1 million interactions per cell, all four methods are capable of distinguishing replicates from non-replicates (Figure 2C), with the best separation provided by HiCRep. This difference may arise because HiCRep explicitly incorporates an explicit smoothing procedure; in contrast, GenomeDISCO uses an implicit smoothing procedure and the other two methods do not perform smoothing at all. Note that these results do not fully resolve question of whether the reproducibility measures will generalize to single-cell data, because in addition to higher sparsity, the variance and noise characteristics of single-cell data is expected to markedly differ from that of bulk Hi-C data. Hence, exploring the applicability of these methods to single-cell Hi-C data more fully is an important direction for future research.

An additional direction for future research is the development of alternative score functions that are designed to focus on particular aspects of chromatin architecture. For example, in the context of single-cell Hi-C analysis, measures have been developed that focus entirely on the genomic distance effect, for use in segregating cells according to cell cycle stages [42]. Similarly, for bulk or single-cell Hi-C, researchers may wish to separately assess whether two cells or cell types exhibit similar chromosome territories, compartment structure, domain structure, or patterns of looping interactions. Developing scores that separately assess these aspects of genome 3D architecture will facilitate automated inference from growing Hi-C data sets.

We release a software package that incorporates the four reproducibility measures and the QuASAR-QC measure (https://github.com/kundajelab/3DChromatin_ReplicateQC). Until recently, proven measures have been lacking and currently there is no standard for measuring for quality and reproducibility of Hi-C data. This tool will both greatly simplify the task of measuring both the quality and reproducibility of Hi-C datasets robustly by using the methods we show to be accurate in this study. We also propose a set of empirical quality and reproducibility thresholds for use at various coverage levels, which are built into the software package to make it easy to determine whether samples pass quality and reproducibility standards (Supp. Table 2).

While the methods we compared are tailored for Hi-C data, similar chromosome conformation capture assays such capture Hi-C [44] and ChIA-PET [45] are used to study three dimensional interactions in the genome. These assays differ from Hi-C due to their targeted nature; however, they share many properties of Hi-C assay, such as the genomic distance effect, and can be represented as a contact matrix similar to Hi-C [46,47]. Reproducibility and quality measures of these assays are lacking in general, raising the possibility of adaptation of the methods we evaluate here to these assays.

In summary, we show that recently proposed Hi-C quality and reproducibility measures accurately measure these qualities on a large collection of real and simulated data. By profiling various parameters of Hi-C contact matrices, we describe best practices for applying and interpreting these measures. We also make available a convenient software tool that simplifies the application of these measures to Hi-C datasets. We hope that adoption of this standard toolkit will help to improve the quality and reproducibility of Hi-C data generated in the future.

## Methods

### Measures of reproducibility

#### HiCRep

This method assesses reproducibility by taking into account two dominant spatial features of Hi-C data: distance dependence and domain structure. The method first smooths the given Hi-C matrices to help capture domain structures and reduce stochastic noise due to insufficient sampling. It then addresses the distance-dependence effect by stratifying Hi-C data according to genomic distance. Specifically, the method consists of two stages.

In the first stage, HiCRep smooths the Hi-C raw contact map using a 2D mean filter, which replaces the read count of each contact with the average counts of all contacts in its neighborhood. The neighborhood size is obtained from a deeply sequenced benchmark dataset using a training procedure. In this analysis, neighborhood size parameter of 20, 5, and 1 are used for the resolutions of 10 kb, 40 kb, and 500 kb, respectively. Smoothing improves the contiguity of regions with elevated interaction, consequently enhancing the domain structures.

In the second stage, HiCRep takes into account the distance dependence effect by a stratification and aggregation strategy. This stage consists of two steps. The algorithm first stratifies the contacts according to the genomic distances of the contacting loci and computes the correlation coefficients within each stratum. HiCRep then assesses the reproducibility of the Hi-C matrix by applying a novel stratum-adjusted correlation coefficient statistic (SCC) to aggregate the stratum-specific correlation coefficients using a weighted average, with the weights derived from the Cochran-Mantel-Haenszel (CMH) statistic. The SCC has a range of [-1, 1] and is interpreted in a way similar to the standard correlation coefficient.

#### GenomeDISCO

This method focuses on two key aspects of contact maps: the need for smoothing, and the multiscale nature of these maps. The need for smoothing arises because contact maps are insufficiently sampled, especially at low sequencing depths. This means that a pair of genomic regions can exhibit a low count either from a lack of contact or from insufficient sampling. This problem is addressed by smoothing the data, essentially assuming that two contact maps are reproducible as long as they capture similar higher order structures, even if they differ in terms of individual contacts. GenomeDISCO investigates contact maps at multiple scales by comparing them at different levels of smoothing and computing a reproducibility score that takes all these comparisons into account.

The smoothing approach is based on random walks on networks. Each contact map is treated as a network, where each node is a genomic region and each edge is weighted by the Hi-C count matrix, following normalization. In this work, square root was used normalization, but similar results were obtained by using alternative normalization methods, including simple row‐ and column-based normalization or Knight-Ruiz normalization [48] (Data not shown). Random walk are performed on networks to smooth the data, asking for each pair of nodes what is the probability of reaching node i from node j, if t steps are allowed in a random walk biased by the edge weights. The smoothed data can be computed by raising the adjacency matrix of our weighted network to the power t. Lower values of t perform local smoothing of the data, revealing structures such as domains, while larger values of t emphasize compartments. This graph-based smoothing scheme aims to preserve sharp domain boundaries that 2D methods may dilute.

To obtain the GenomeDISCO reproducibility score, each contact map is separately smoothed across a range of t values. For each value of t, the L1 distance (i.e., the sum of the absolute values in the difference matrix) between the two smoothed contact maps is computed and normalized by the average number of nodes with nonzero total counts across the two original contact maps compared. Afterwards, a combined distance between the two contact maps is obtained by computing the area under the curve of the L1 difference as a function of t. This allows us to consider multiple levels of smoothing and thus multiple scales when computing our scores. Finally, this distance is converted into a reproducibility score as follows:

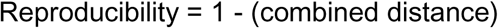

This score is in the range [-1,1], with higher scores representing higher reproducibility. This is because, for each node, the maximum L1 difference is 2, corresponding to the case when the node has mutually exclusive contacts in the two contact maps being compared. Thus, the combined distance lies in the range [0, 2], making the reproducibility score fall in the range [-1, 1].

Parameter optimization on an orthogonal dataset revealed the optimal t={3} [34], which was used in this study.

In all pairwise comparisons in this paper, the sample with higher coverage was downsampled to match the coverage of the other sample.

#### HiC-Spector

The starting point of spectral analysis is the Laplacian matrix L, which is defined as L=D-W, where W is a symmetric and non-negative matrix representing a chromosomal contact map, and D is a diagonal matrix in which *D*_*ii*_ = Σ_*j*_*W*_*ij*_. The matrix L is further normalized by the transformation *D*^−1/2^*LD*^−1/2^, and its leading eigenvectors are found. As in other commonly used dimensionality reduction procedures, the first few eigenvalues are of particular importance because they capture the basic structure of the matrix, whereas the latter eigenvalues are essentially noise. Given two contact maps *W*^*A*^ and *W*^*B*^, their corresponding Laplacian matrices *L*^*A*^ and *L*^*B*^ and corresponding eigenvectors are calculated. Let 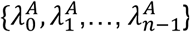, and 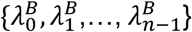 be the spectra of *L*^*A*^ and *L*^*B*^ and 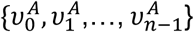 and 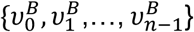 be their normalized eigenvectors. A distance metric is defined as:

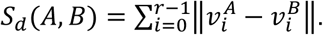

Here || || represents the Euclidean distance between the two vectors. The parameter r is the number of leading eigenvectors used. In general, *S*_*d*_ provides a metric to gauge the similarity between two contact maps. The distance is then linearly rescaled to a reproducibility score ranging from 0 to 1.

#### QuASAR-Rep

The Quality Assessment of Spatial Arrangement Reproducibility (QuASAR) measure uses the concept that within a distance matrix, as the distance between two features approaches zero, the correlation between the rows corresponding to those two features approaches one. This relationship is exploited by calculating the interaction correlation matrix, weighted by interaction enrichment. To determine reproducibility across replicates, the correlation of weighted correlation matrices is calculated as follows. In every case, matrices are first filtered by removing intra-chromosomal interaction matrix rows and columns such that all remaining rows and columns contain at least one non-zero entry within 100 bins up‐ or downstream of the diagonal. The background signal-distance relationship is estimated as the mean number of reads for each inter-bin distance. The interaction correlation matrix is calculated across all pairwise sets of rows and columns within 100 bins of each other from the log-transformed enrichment matrix (non-filtered counts divided by background signal-distance values), excluding bins falling on the diagonal in either set. For a given pair of rows A and B, the correlation is calculated from all columns within 100 bins of both A and B, excluding filtered columns. The interaction matrix is then found by adding 1 to valid entries and taking the square root. The weighted correlation matrix is an element-wise product of the correlation matrix and the interaction matrix divided by the sum of all valid interaction matrix entries. The replication score is the correlation of weighted correlation matrices between two samples. Note that, to distinguish the use of QuASAR for assessing reproducibility versus data quality (described below), we refer in the main text to “QuASAR-Rep” and “QuASAR-QC”.

#### Processing of reproducibility scores

All the reproducibility measures we use in this study assign a reproducibility score to a pair of Hi-C contact matrices. Due to the sparsity and noise nature of inter-chromosomal matrices, reproducibility scores are only calculated for intra-chromosomal matrices. The final reproducibility score assigned to a pair of Hi-C experiments in this study is the mean of the reproducibility scores assigned to pairs of Hi-C contact matrices of each chromosome.

#### Empirical reproducibility score thresholds

To infer empirical thresholds for distinguishing non-replicates for biological replicate pairs for each method, we used the distribution of reproducibility scores assigned to non-replicate pairs and biological replicate pair at a given coverage level. Similar to the concept of a maximal margin hyperplane, the empirical threshold we inferred is midpoint of the reproducibility score of the highest scoring non-replicate pair and the reproducibility score of the lowest scoring biological-replicate pair. For each coverage level from 30 million Hi-C interactions to 5 million interactions, we inferred a single empirical threshold for each reproducibility metric. These thresholds are available in Supp. Table 2.

### Measures of quality

#### QuASAR-QC

The sample quality measure from QuASAR (“QuASAR-QC”) uses the same transformation as described above for reproducibility. However, instead of looking at weighted correlation matrices between samples, the quality score is found by taking the weighted correlation mean across all chromosomes and then subtracting the unweighted correlation mean across all chromosomes.

#### TAD boundary calling and analysis

TAD boundaries were identified using the insulation score [40]. This score captures the density of signal in the Hi-C contact matrix around the diagonal, as a function of genomic position. Because the signal is weaker at the boundary of two TADs, minima in the insulation score profile correspond to TAD boundaries. We used the TAD calling software described in Giorgetti et al. [5], employing the previously used parameters (‐‐ss 80000 ‐‐im iqrMean ‐‐is 480000 ‐‐ids 320000) for calculation of the insulation score and identification of minima.

To characterize the effects of noise and coverage on TAD boundary identification, we used noise injected and downsampled datasets as explained before and used insulation score method as desribed in the previous section. For noise injected datasets, we found that number of identified TADs across the genome are only altered at the highest noise levels: the number of total TADs increased only by 5% with 50% noise injection (Supp. Fig. 11). Consistent with the changes in the total number of TADs, the distribution of TAD sizes is only altered at high noise levels. For 7 out of 11 cell types, we detect a statistically significant reduction in the TAD size distribution (P < 0.01, Kolmogorov-Smirnov test) only at either 40% or 50% noise (Supp. Fig. 12). Furthermore, positions of TAD boundaries are not altered with increasing noise levels. For 11 cell types, we calculated the distances between the TAD boundaries of the combined noise-free biological replicate and the TAD boundaries from noise injected replicates. These distances were compared against the TAD boundary distances from biological replicate pairs, which serves as a baseline for how much the TAD boundaries fluctuate between different replicates from the same cell type. Again, we found that the boundary distances are significantly larger than the baseline distribution (one sided KS test, P < 0.05) only at the 50% noise level for four cell types and never larger for the remaining four cell types (Fig. 4C, Supp. Fig. 13).

We adopted the same approach for investigating the effect on coverage on insulation score identified TADs. For each of the six downsampled cell lines, we identified TADs using insulation score method and compared total number of TADs, the size distribution of TADs and the differences between TAD boundaries between the original replicate and downsampled replicates. We observe that the total number of TADs detected and TAD size distributions are similar at all coverage levels (Supp. Fig. 16 and 17). We calculated the distances between TAD boundaries identified from downsampled replicates against the TAD boundaries from original biological replicates, and we compared this distribution against the distances between biological replicates as a baseline. For five of the six cell types, downsampling causes the TAD boundaries to shift away from the original boundaries significantly (Kolmogorov-Smirnov test, P <0.05) only 10 million and lower number of interactions, further supporting the idea that TAD boundary by insulation score detection is mostly robust to low coverage (Fig. 4F, Supp. Fig. 18).

#### Number of significant contacts

For a given normalized Hi-C contact map, we computed the number of contacts that are deemed statistically significant using Fit-Hi-C [38]. Hi-C contact maps were binned at 40kb resolution and normalized using the Knight-Ruiz matrix balancing algorithm [42]. Deeply sequenced Hi-C data from 2 cell types were binned at 10kb and 40kb resolutions for Fit-Hi-C analysis (Supp. Table 2). Fit-Hi-C assigns a statistical significance to each contact between two bins by assigning a p-value and a q-value. For each experiment, we counted the number of contacts that are above a given q-value threshold for every intra-chromosomal interactions and aggregated them over all chromosomes and used this sum as the total number of significant contacts for a given experiment.

### Mapping statistics

We have used three statistics to summarize alignment quality, valid Hi-C fragment pairs, and the ratio of intrachromosomal and interchromosomal Hi-C interactions. A thorough description of these statistics and their application is reviewed in Lajoie et al [24]. First statistic we use is percentage of aligned pairs, which corresponds to the percentage of Hi-C fragment pairs that uniquely map to the genome on both sides. Typically, single sided and non-unique alignments are discarded in Hi-C pipelines [23,24]. The second statistic is invalid pairs, which the percentage of aligned pairs that map against the same restriction fragment. These fragment pairs are non-informative since they do not correspond to a fragment between two different regions [24]. The third statistic is the percentage of intra-chromosomal valid pairs. Random ligations are much more likely to result in interchromosomal fragments; thus a high ratio of non-informative random ligation events result in an enrichment of inter-chromosomal interactions and a depletion of intra-chromosomal interactions [24]. The fourth statistic is the percentage of PCR duplicates, which is estimated from the number of aligned pairs that map to the exact same coordinates as another aligned pair [24].

### Simulation of noisy Hi-C matrices

To generate noise for Hi-C data in a realistic manner, we simulated two Hi-C contact matrices that would result from two processes that are not dictated by the 3D organization of chromatin. These “pure noise” matrices are mixed with the real Hi-C contact matrix to generate the final, noisy Hi-C matrix. The first noise matrix models the genomic distance effect, namely the higher probability of observing a Hi-C interaction between two regions that are close along the one dimensional length of a chromosome. Because such regions are constrained to be close to each other, they are more likely to interact compared to more distal regions, in the absence of any higher order structure. This effect has been documented early on and is generally corrected in Hi-C contact matrices to better visualize medium and long range interactions [1]. The second noise matrix models the ligation of non-crosslinked DNA fragments during the ligation step of the Hi-C protocol. Fragments pairs that results from random ligation are uninformative since they can link two regions independently of 3D organization.

Additionally, the Hi-C assay is subject to the same biases that other next generation sequencing assays suffer from. These biases result include a bias in favor of GC rich regions and a bias against regions of low mappability. During the generation of both types of noise matrices, we factored in such biases by using the sum of each row as a proxy for the the overall bias of a bin. Coverage normalization of Hi-C matrices [1] similarly uses marginals to counter such biases.

To generate the genomic distance noise matrix G, we sampled from empirical distributions derived from of real Hi-C matrix. In this setting, the genomic distance D is defined as the number of bins that lie between a pair of bins i and k, i.e. |*i*-*k*| = *D*. For every value of D, we build a vector S by collecting the set of real Hi-C matrix entries M_ik_ for which |*i*-*k*| = *D*. We then randomly select values from S for insertion into G, again considering only entries G_ik_ for which |*i*-*k*| = *D*. This sampling strategy effectively shuffles the matrix entries in M at a fixed distance, thus preserving the original genomic distance effect while disrupting other higher order structures. However, instead of uniformly sampling from S, we adopted a stratified sampling strategy to better model GC and mappability biases. Specifically, S was broken into multiple strata before sampling. The strata are determined by products of marginals, i.e. M_ik_ is assigned to a certain stratum based on the product of the marginals of bin i and bin k. For a given value of D, we chose stratum size in such a way that each each stratum contains 100 elements. When sampling the G_ik_, we sampled a value from from the stratum that Mik belongs to. By repeating the stratified sampling for every value of D, the final matrix G is obtained.

To generate the random ligation noise matrix R, we generated random Hi-C interactions and aggregated them to build a Hi-C contact matrix. We generated these interactions by randomly choosing two bins *i* and *k*, and adding one to the matrix entry R_ik_ in the random noise contact matrix. Instead of sampling the bins uniformly, the probability of sampling a bin was set proportional to marginal of that bin, thus modeling the GC and mappability bias of each bin. The sampling process was repeated N times, where N is the total number of interactions in the original Hi-C contact matrix M, to generate a random ligation noise matrix.

After both noise matrices are generated from the original Hi-C matrix, these matrices were mixed in varying proportions to generate a series of noisy Hi-C matrices. Each such matrix is a mixture of three matrices: a real matrix, a genomic distance noise matrix, and a random ligation noise matrix. To generate a simulated matrix with *c* total counts from, we sampled counts uniformly at random from one real and two simulated matrices at a given target ratios. In practice, we varied the total proportion of noise from 0% to X%, and for each total noise level we consider two settings for the relative proportions of genomic distance noise random ligation noise: we either used one third of matrix G and two thirds of matrix R, or vice versa. We note that most analyses in this study were robust to either scenario.

The software for injecting noise into Hi-C contact matrices is available at https://github.com/gurkanyardimci/hic-noise-simulator.

### Downsampling

Downsampled data sets were generated by converting an input Hi-C matrix into a set of pairwise individual intra-chromosomal interactions and uniformly sampling a given number of interactions from this set. Following downsampling, we re-binned the set of chosen interactions into a Hi-C matrix.

For analysis of reproducibility measures, we limited the analysis to real data from six cell types with replicates of at least 30 million interactions, and we downsampled each individual replicate to have a wide range of total interactions (30 × 10^6^, 25 × 10^6^, 20 × 10^6^, 15 × 10^6^, 10 × 10^6^, 5 × 10^6^, 10^6^). Using a single pseudo-replicate and a single biological replicate pair for each cell type and 15 non-replicates at each coverage level, we generated a total of 189 replicate pairs. These datasets were used for testing the ability of each method to distinguish among different replicate types at lower coverage levels, and for explicitly profiling the dependence of reproducibility scores on coverage levels.

For analysis of QC measures, we generated downsampled biological replicates from the same six cell types to have fewer interactions (30 × 10^6^, 25 × 10^6^, 20 × 10^6^, 15 × 10^6^, 10 × 10^6^, 5 × 10^6^, 10^6^), resulting in a set of 84 matrices. In addition, we applied the same setup to deeply sequenced datasets from two cell types at a wider range of coverage values (30 × 10^6^, 60 × 10^6^, 120 × 10^6^, 240 × 10^6^, 400 × 10^6^), at multiple resolutions, resulting in 30 matrices. For each downsampled matrix, we calculated QuASAR scores and identified statistically significant long range contacts and TAD boundaries.

### Generation of pseudo-replicates

Given two biological replicate experiments, we generated pseudo-replicates by aggregating the two replicates and downsampling from the combined matrix. Combination of two biological replicates is performed by summing the two Hi-C contact matrices of these replicates. Following combination, the resulting combined Hi-C matrix is downsampled as described above to generate pseudo-replicates. We forced the pseudo-replicates to have the average of total number of interactions of two seed biological replicates.

### Resolution analysis

To investigate whether an optimum resolution exists for a given sample, we profiled the reproducibility scores assigned to biological replicate pairs from four cell types: A549, G410, LNCaP and HepG2. Hi-C data from the first three cell types were generated by the HindIII restriction enzyme, whereas the HepG2 data was generated using DpnII. These samples also exhibit differing levels of coverage (Supp. Table 1). In this analysis, we binned the contact matrix of each replicate at 40kb, 20kb, 10kb and 5kb resolution and calculated the various reproducibility scores assigned to each biological replicate pair. For this analysis only, we limited the computation of reproducibility scores to the contact matrices of chr22.

## Availability of Data and Materials

The Hi-C data used for generating simulated hi-c matrices and the evaluation of quality control and reproducibility methods is publicly available at https://www.encodeproject.org/. The accession code of each dataset is available in Supplemental Table 2. The code for running reproducibility and quality controls methods is publicly available at https://github.com/kundajelab/3DChromatin_ReplicateQC. Software for simulating noise injected matrices can be accessed at https://github.com/gurkanyardimci/hic-noise-simulator.

## Authors’ Contributions

GGY, JD, WSN designed the experiments. Y.Z. and B.R.J. generated and processed the data. G.G.Y., H.O., M.E.G.S., O.U., K.K.Y., T.Y, A.C., A.K., F.A. ran the experiments. G.G.Y and W.S.N. analyzed the results. G.G.Y, O.U. and W.S.N made the figures. O.U. wrote the 3DChromatin_ReplicateQC software, with input from M. E. G. S., K. K. Y. and T. Y. All authors contributed to the preparation of the manuscript.

## Acknowledgements

We thank Giancarlo Bonora and Kate Cook for helpful discussions.

---

**Supp. Fig. 1.**
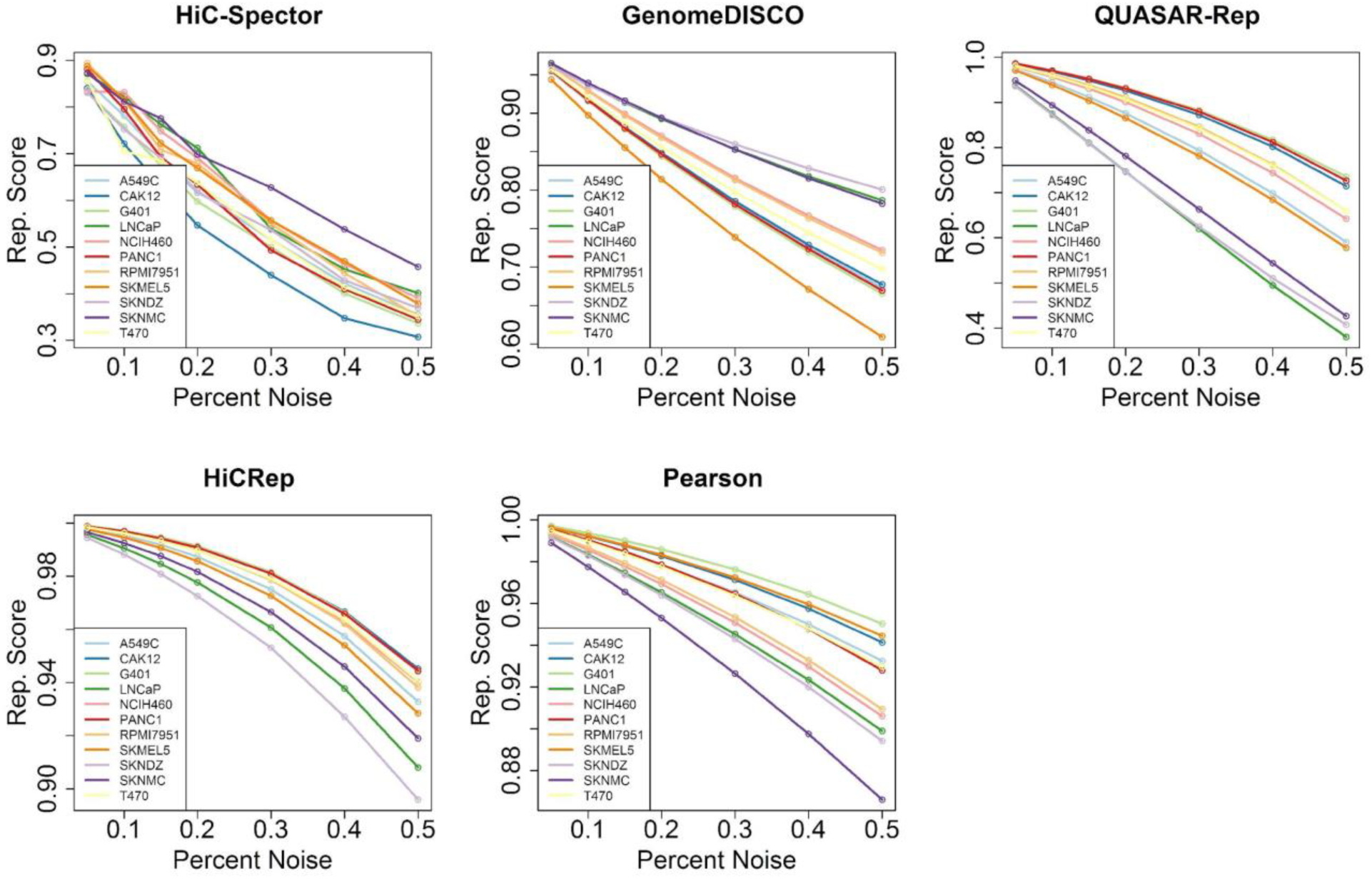
Reproducibility scores assigned to each noise injected replicate pair for each cell type for 33% random ligation configuration. For every cell type and every measure, we see a monotonic trend of decreasing scores with higher levels of noise. The same trends for each measure are observed in the 66% random ligation noise configuration (not shown).

**Supp. Fig. 2.**
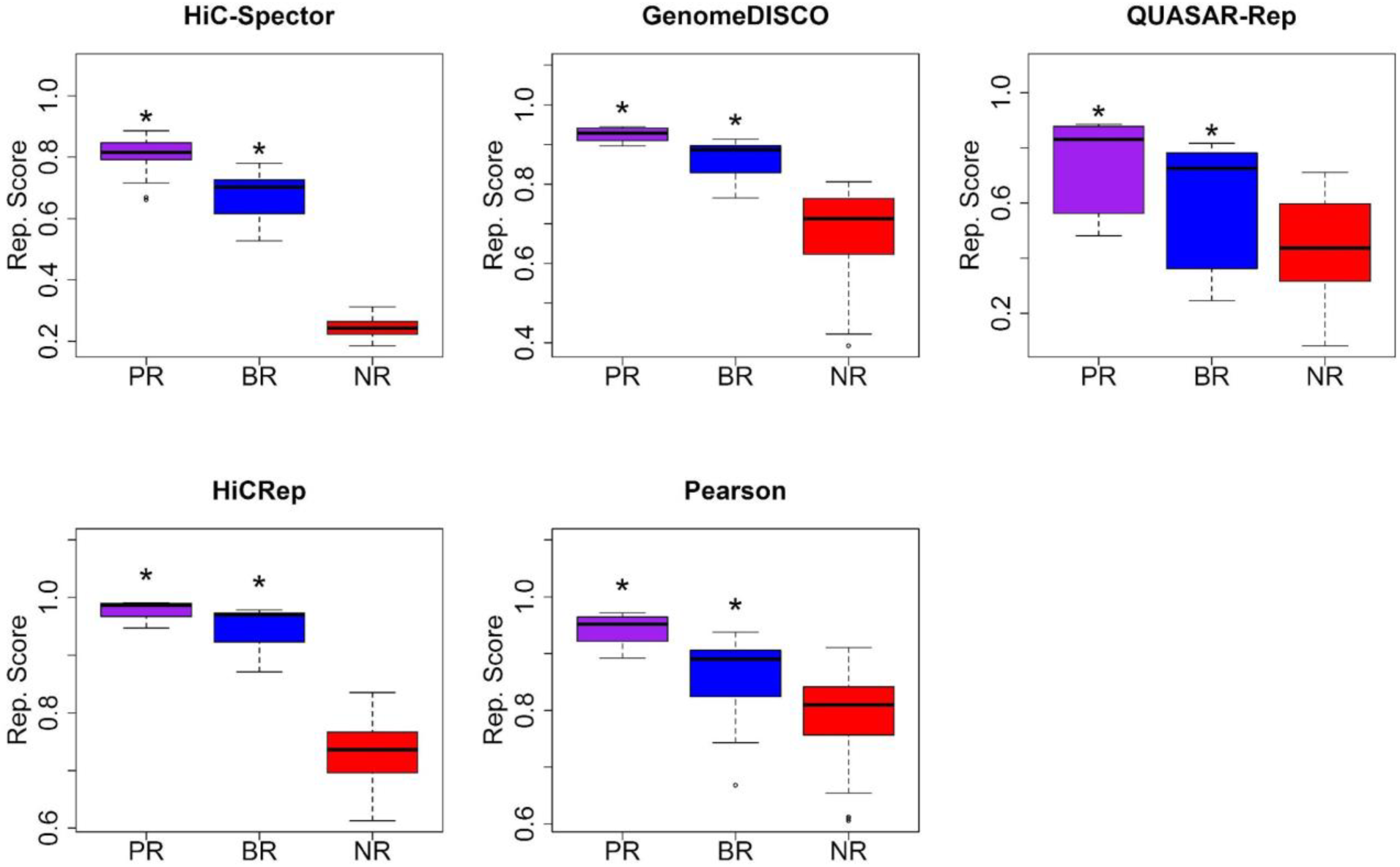
Boxplots showing the distribution of reproducibility scores assigned to each replicate pair category by each measure: pseudo replicates (PR), biological replicates (BR) and non-replicates (NR). Asterisks indicate that the marked distribution is significantly larger than the preceding one according to a one-sided Kolmogorov-Smirnov test (P < 0.01).

**Supp. Fig. 3.**
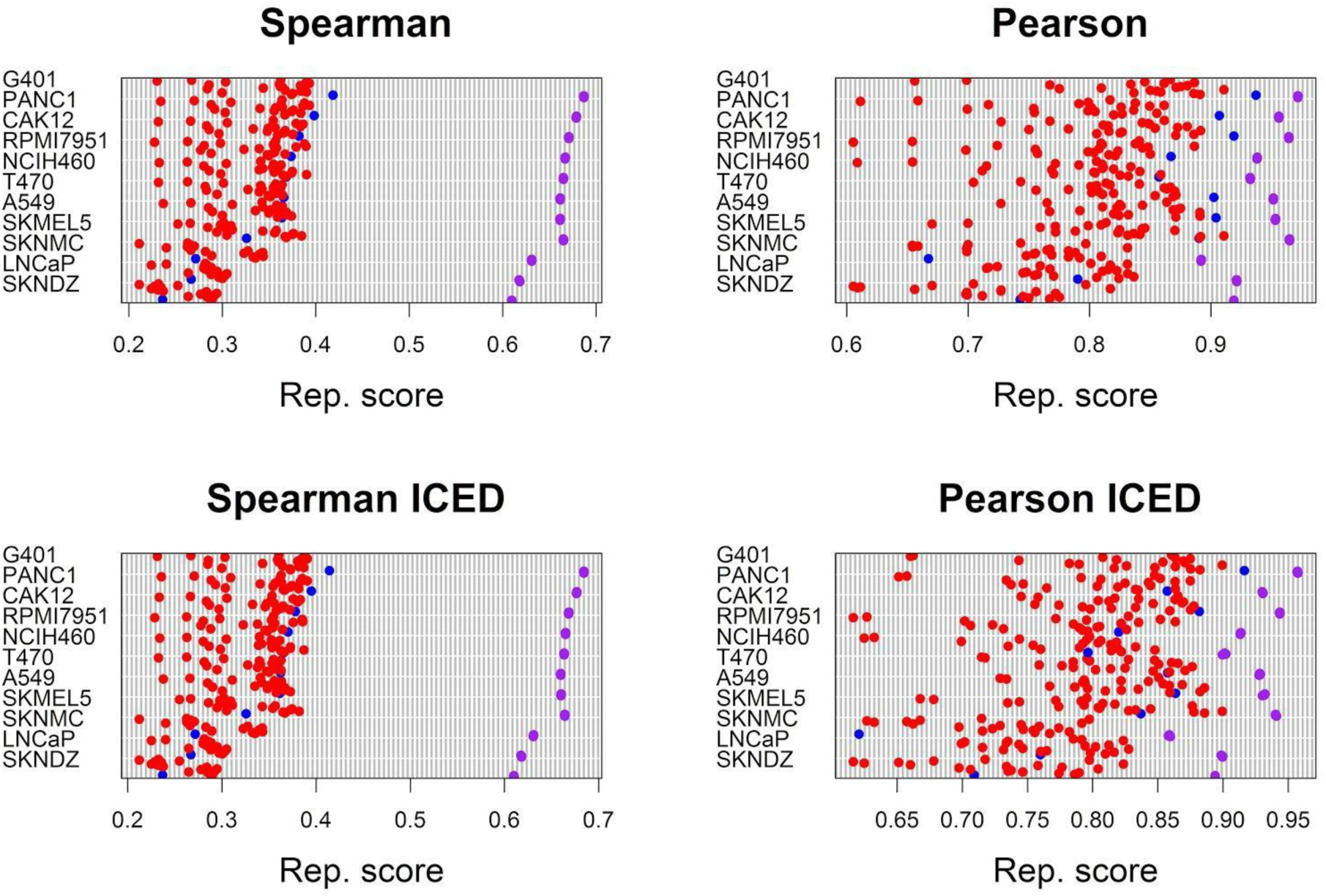
Comparison of Pearson and Spearman correlation coefficients on raw and normalized Hi-C data. Correlation coefficients assigned to biological replicate (blue), non-replicate (red) and pseudo-replicate (purple) pairs are plotted for each cell type. All of these experiments used HindIII digestion.

**Supp. Fig. 4.**
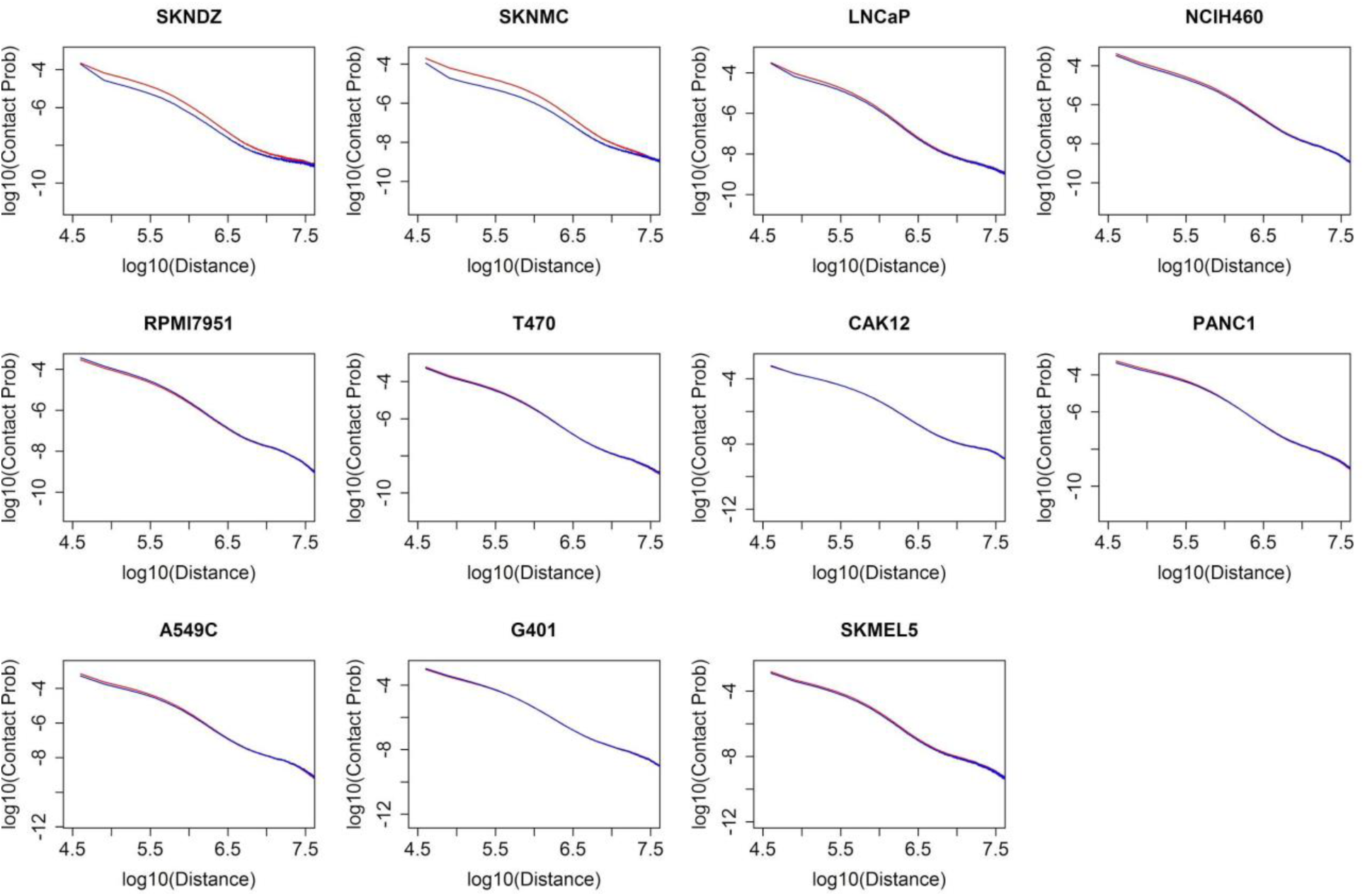
Contact probability curves for biological replicates pairs from each replicate pair. The curve for first replicate is shown in red and the second replicate is in blue.

**Supp. Fig. 5.**
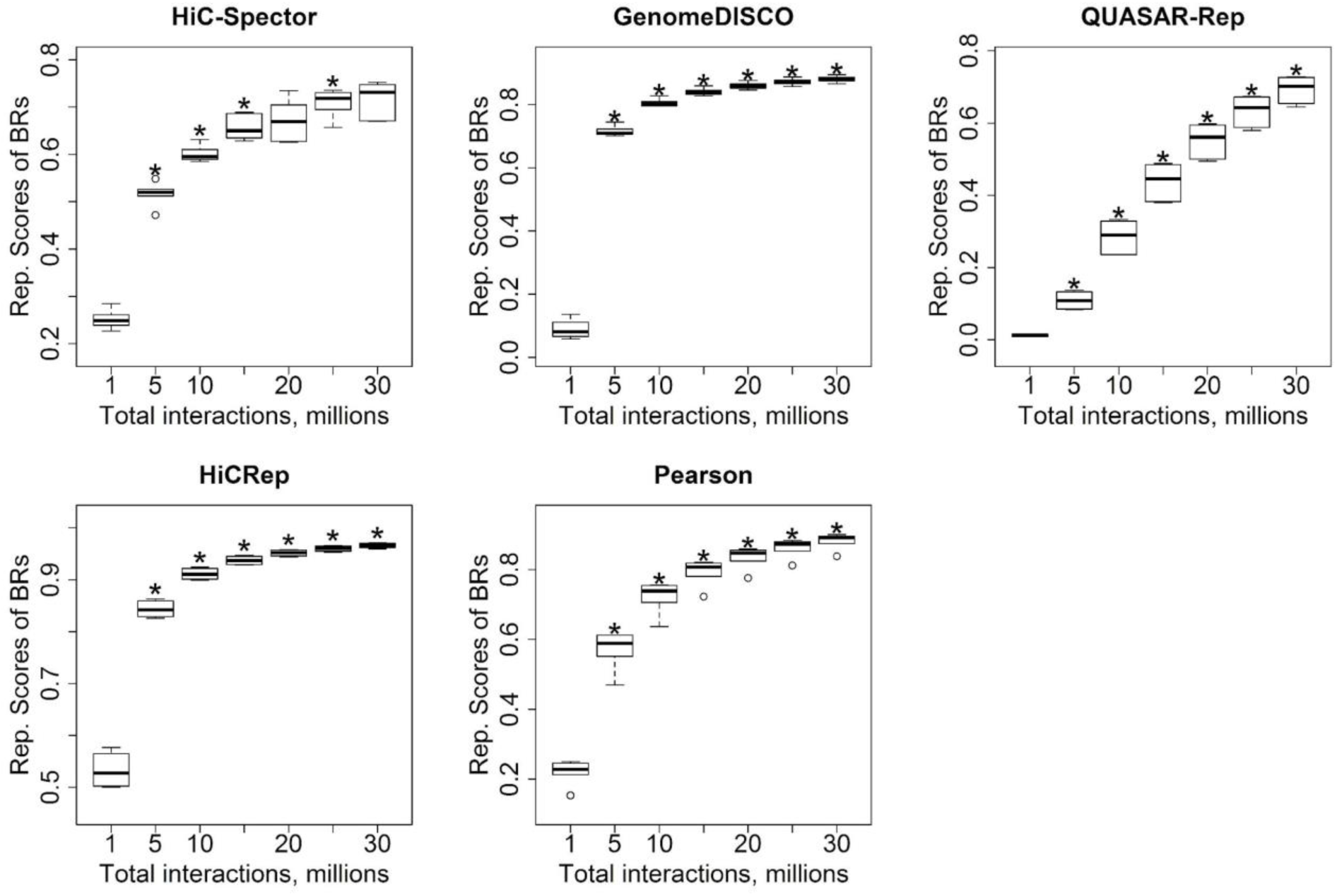
Boxplots showing the distribution of reproducibility scores assigned to six downsampled biological replicates at each coverage level. Asterisks above each distribution indicate that the distribution of reproducibility scores assigned to that distribution is significantly larger than the previous distribution according to a one-sided Wilcoxon signed rank test (P < 0.05)

**Supp. Fig. 6.**
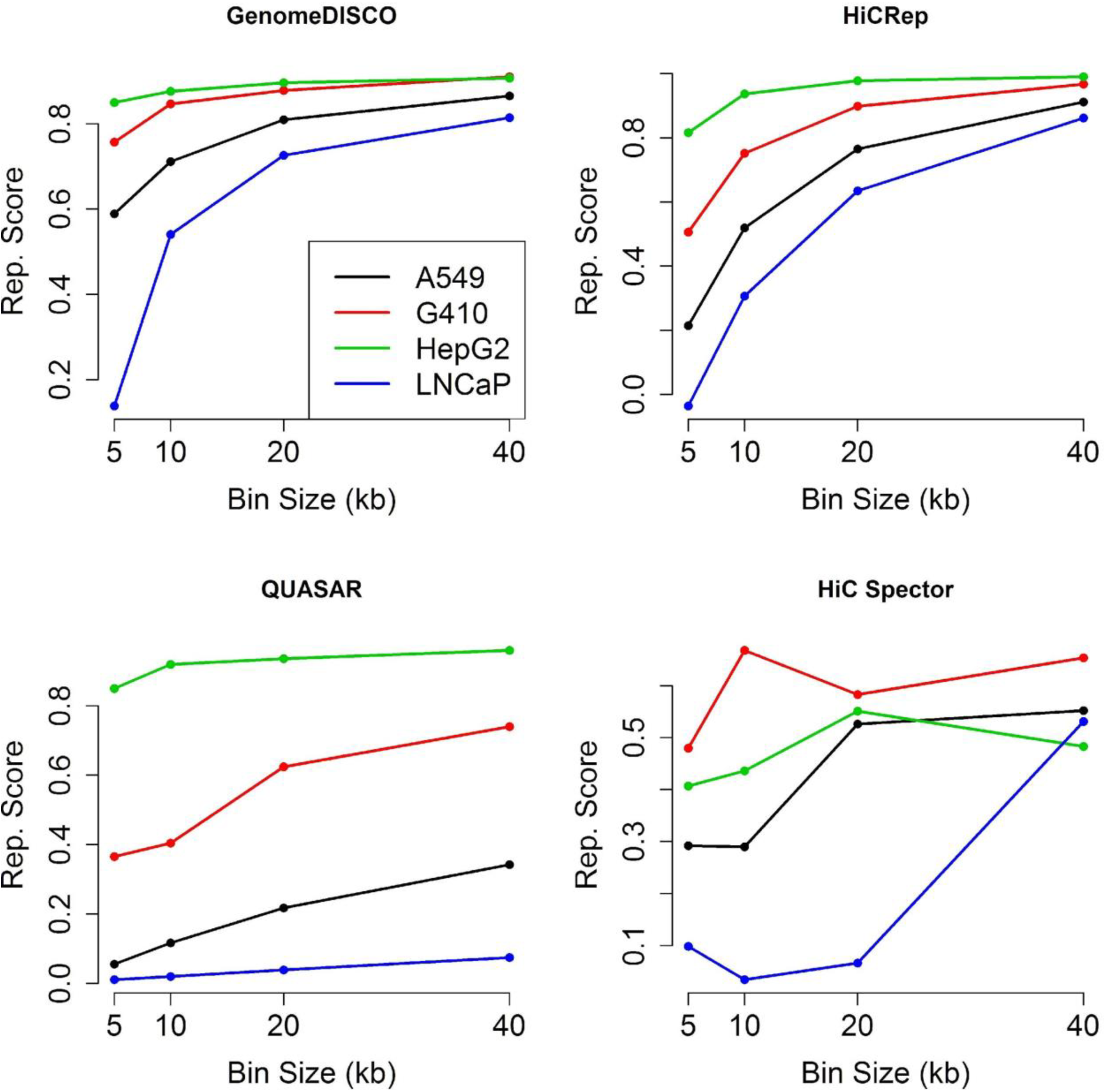
Relationship between reproducibility score and resolutionbin size. Each panel plots a specific reproducibility score as a function of resolution (5, 10, 20, 40 kb). Each of the four series corresponds to a biological replicate from a specified cell line.

**Supp. Fig. 7.**
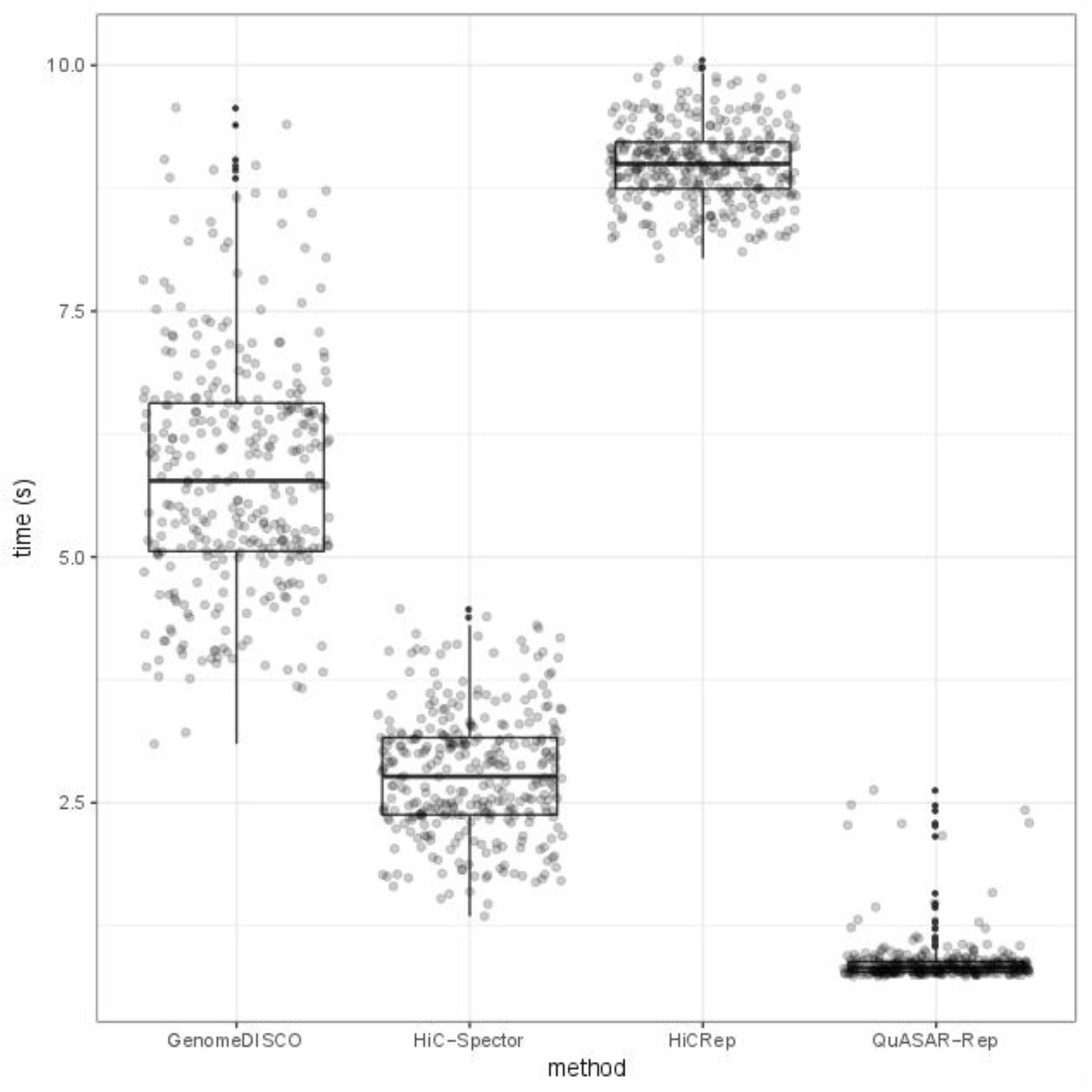
Run times for each reproducibility measure. Each boxplot shows the distribution of run times on 326 replicate pairs for each reproducibility measure. The times we report are wall-clock times, as reported by the “time” command in Unix. All tests were run on an Intel Xeon CPU E5-2683 v3 running at 2.00GHz.

**Supp. Fig. 8.**
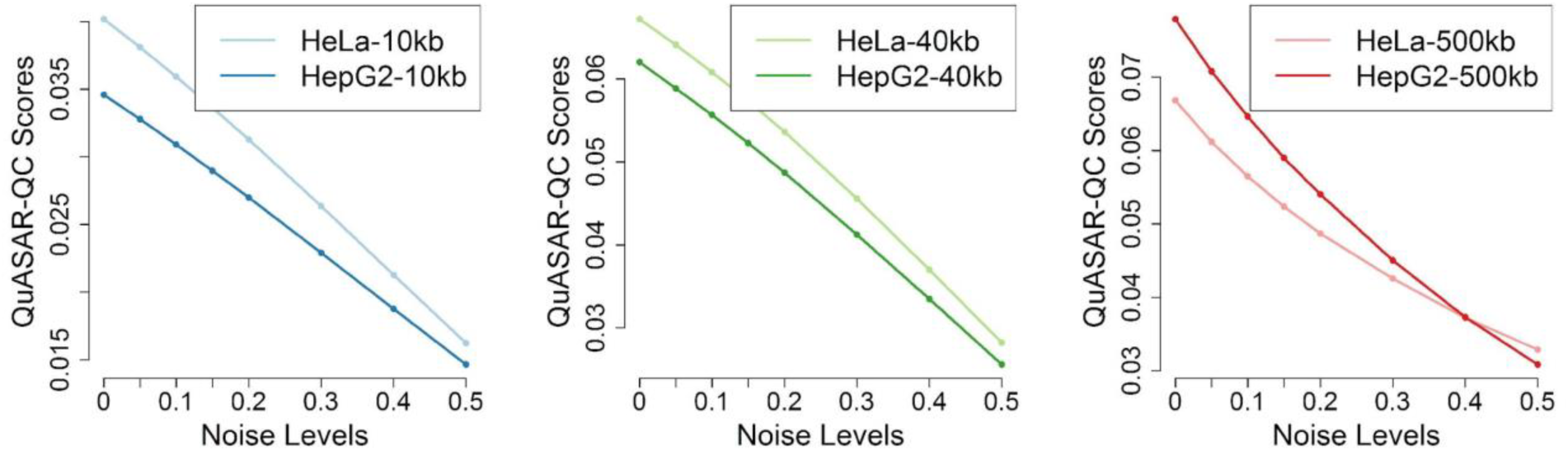
QuASAR-QC scores assigned to noise injected simulated datasets from deeply sequenced replicates. QuASAR-QC scores decrease with increasing levels of noise at all resolutions.

**Supp. Fig. 9.**
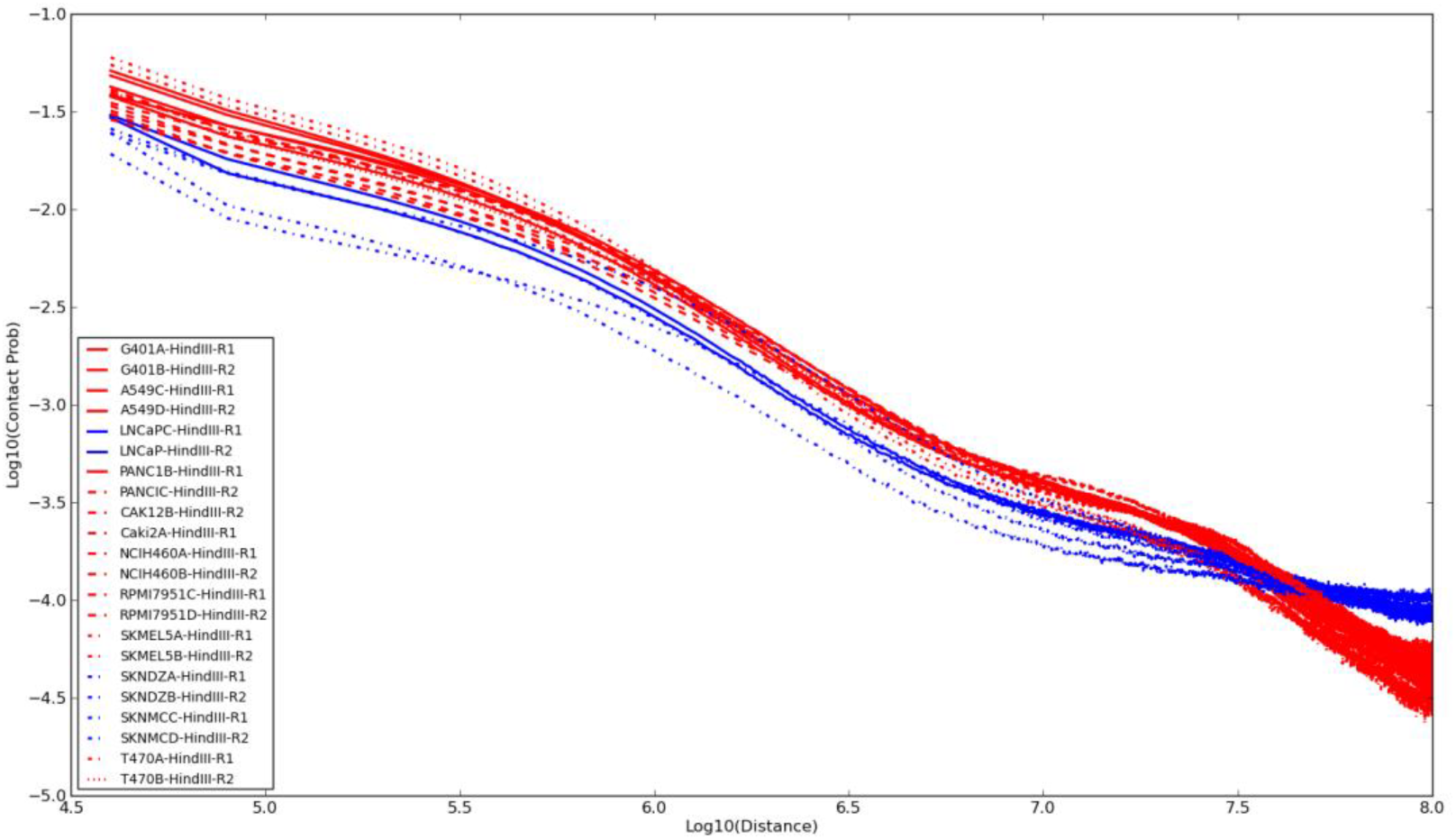
Curves showing the probability of contact between two loci against genomic distance. The blue curves are generated using Hi-C experiments done on SKNDZ, SKNMC and LNCaP cell lines.

**Supp. Fig. 10.**
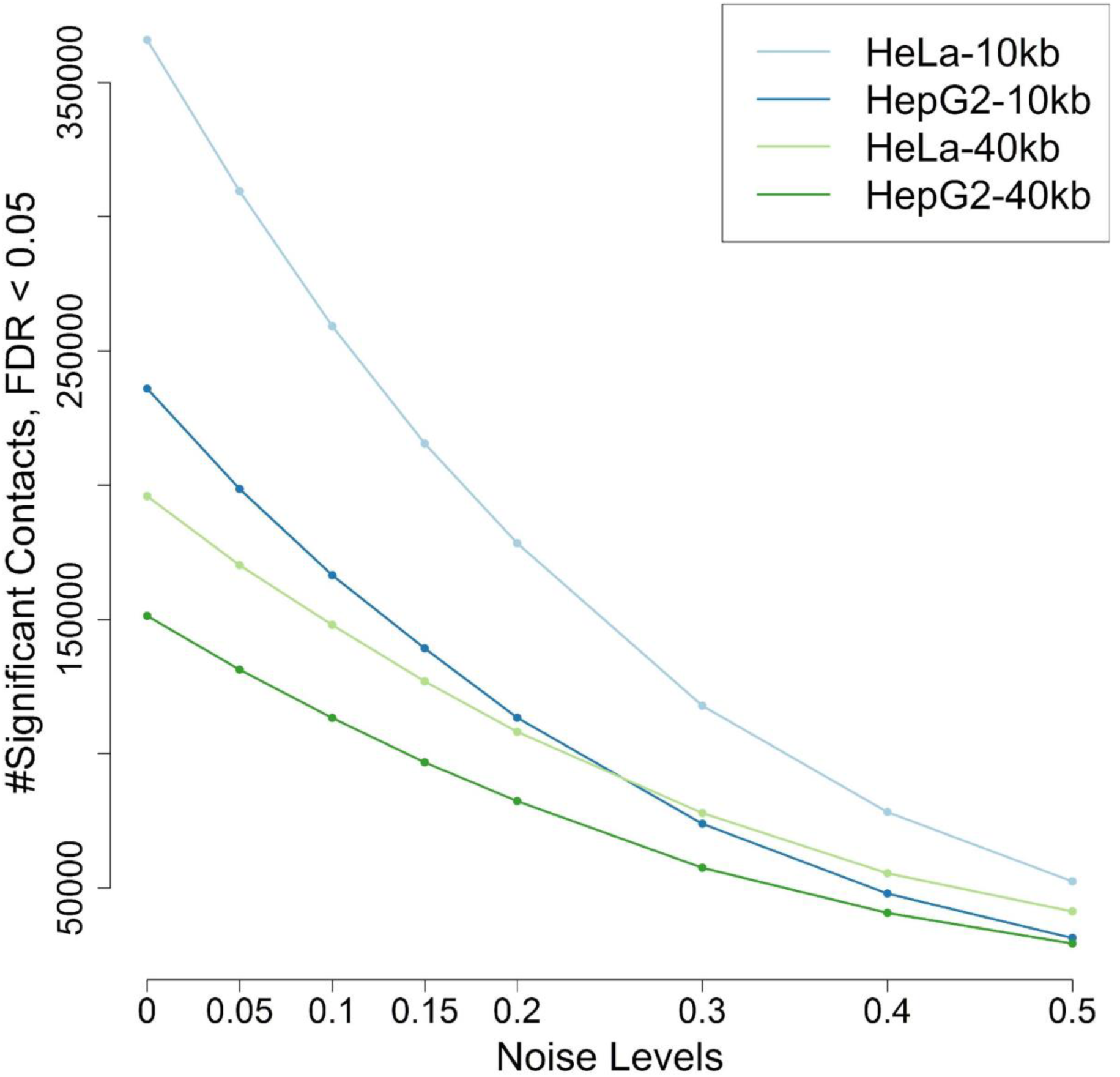
Total number of significant mid-range contacts identified by Fit-Hi-C with an FDR threshold of 0.05 from simulated datasets at 10kb and 40kb resolutions, generated from deeply sequenced datasets. The total number of contacts for each cell type and resolution decreases monotonically with increasing levels of noise.

**Supp. Fig. 11.**
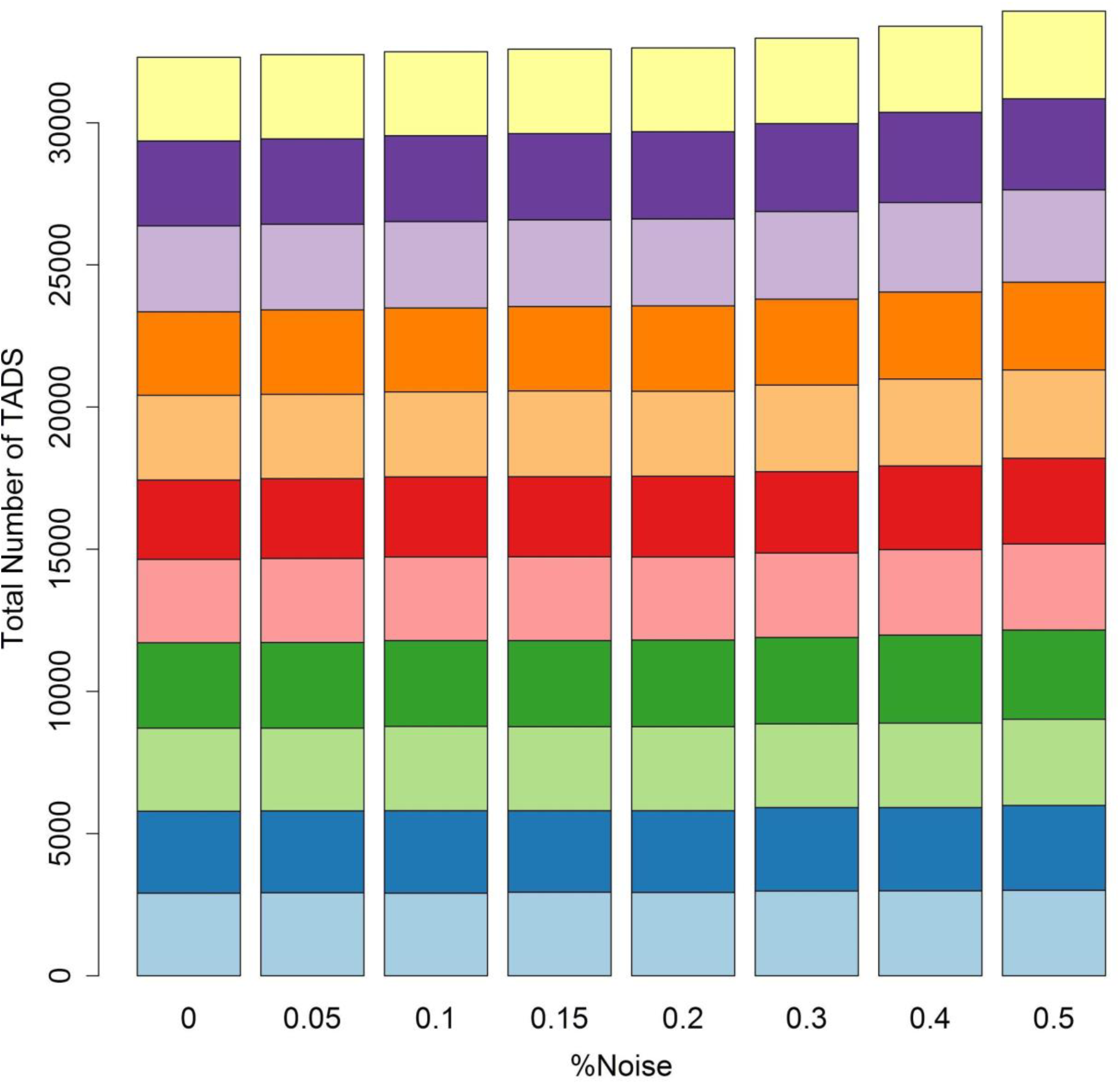
Bar plots showing the number of TADs for each cell line (coded by color) at each noise injection level.

**Supp. Fig. 12.**
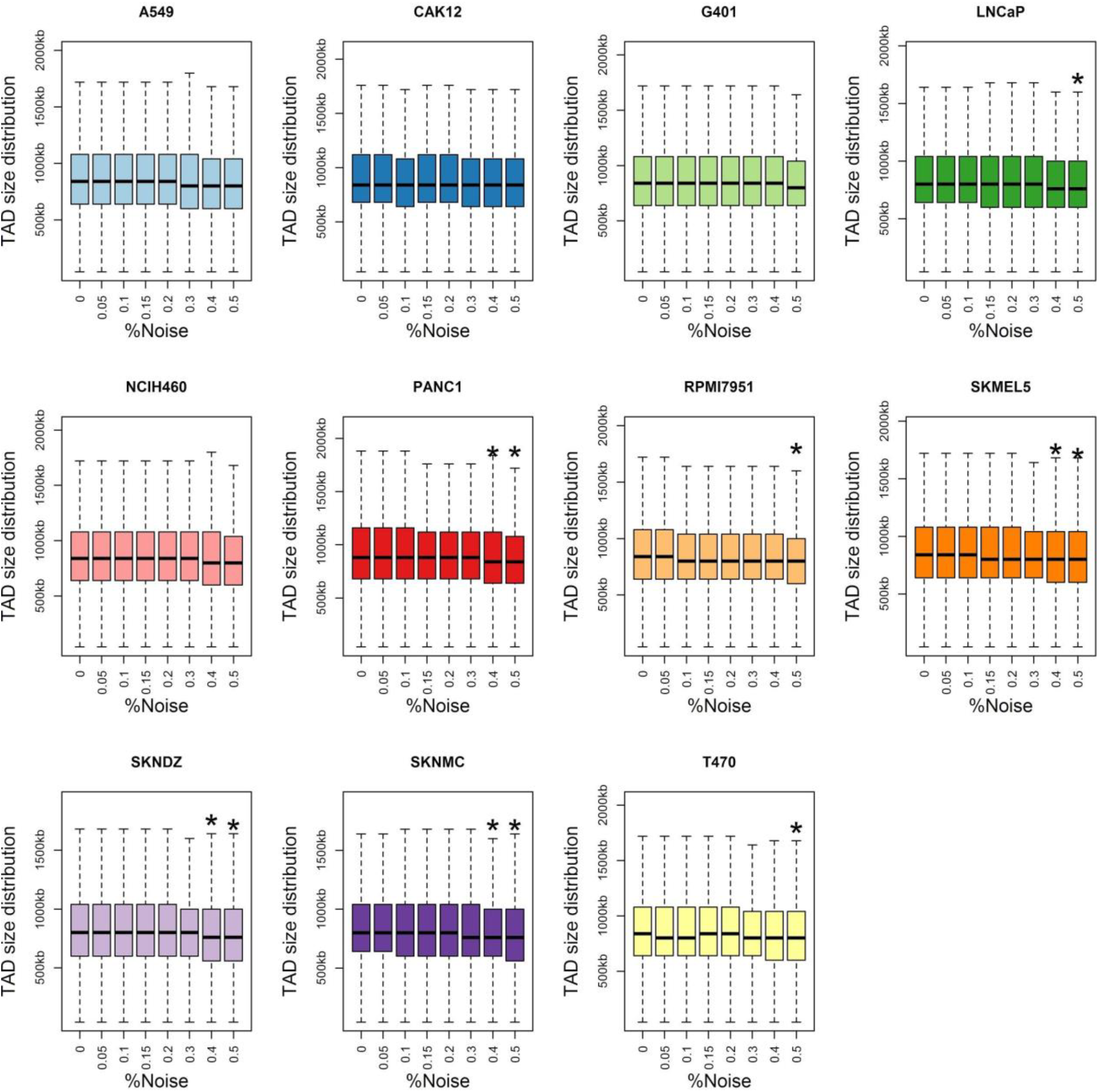
Boxplots showing the distribution of TAD sizes at each noise injection level. Each plot corresponds to a simulated dataset from an individual cell type. Distributions marked with an asterisk are significantly different from the original distribution TAD sizes detected from the noise-free replicate (KS test, P < 0.01).

**Supp. Fig. 13.**
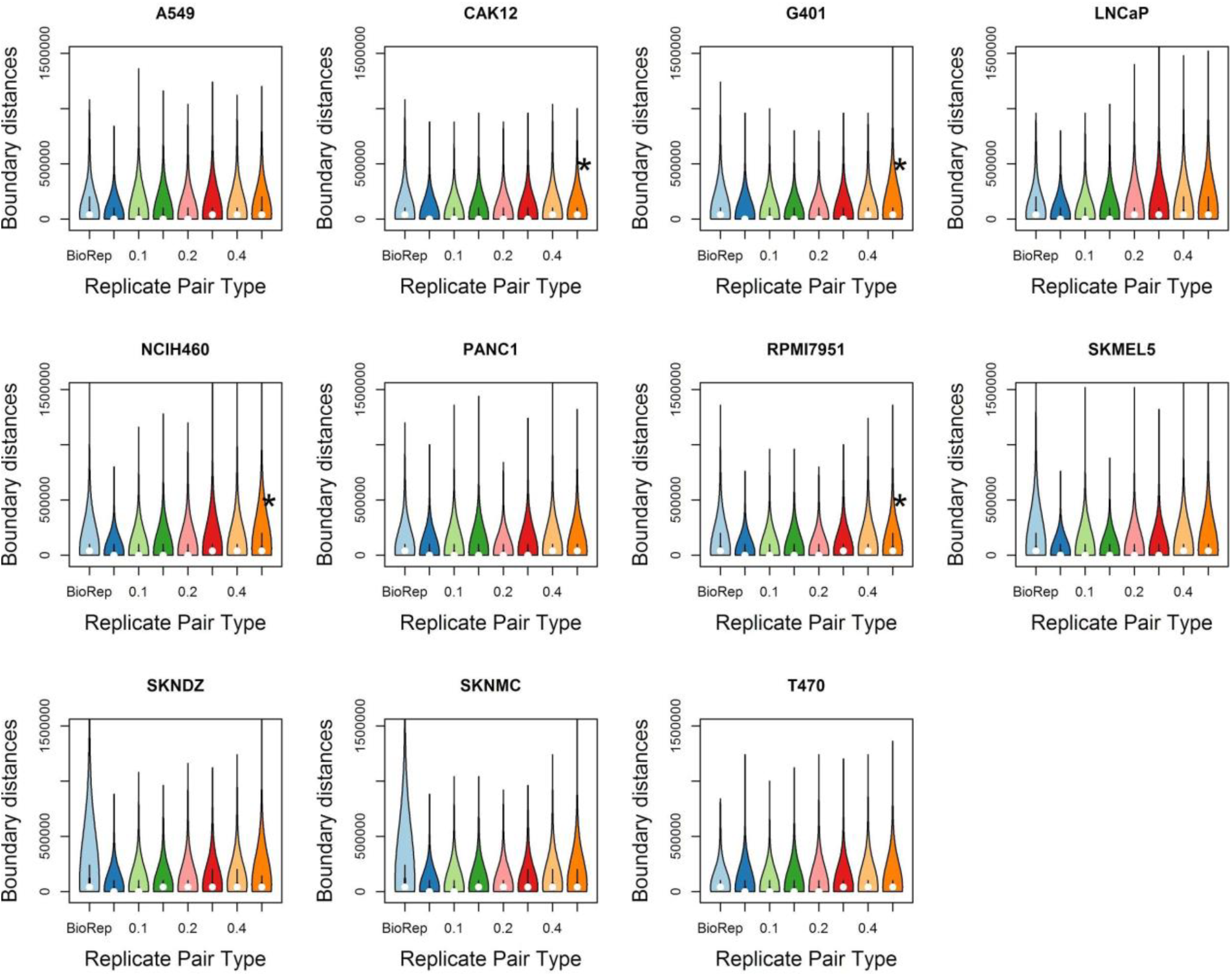
Violin plots showing the distribution of distances between domain boundaries between biological replicates and simulated replicates. Each panel corresponds to a single cell type.

**Supp. Fig. 14.**
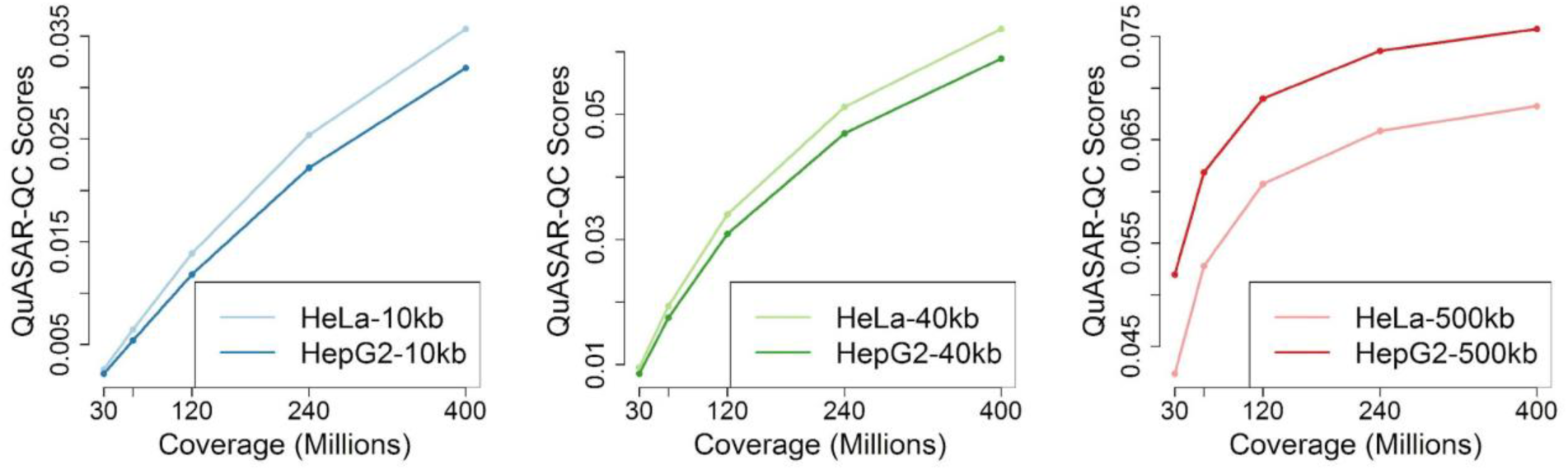
Curves showing the QuASAR scores assigned to deeply sequenced cell types downsampled to 30, 60, 120, 240 and 400 million interactions at 10kb, 40kb and 500kb resolutions.

**Supp. Fig. 15.**
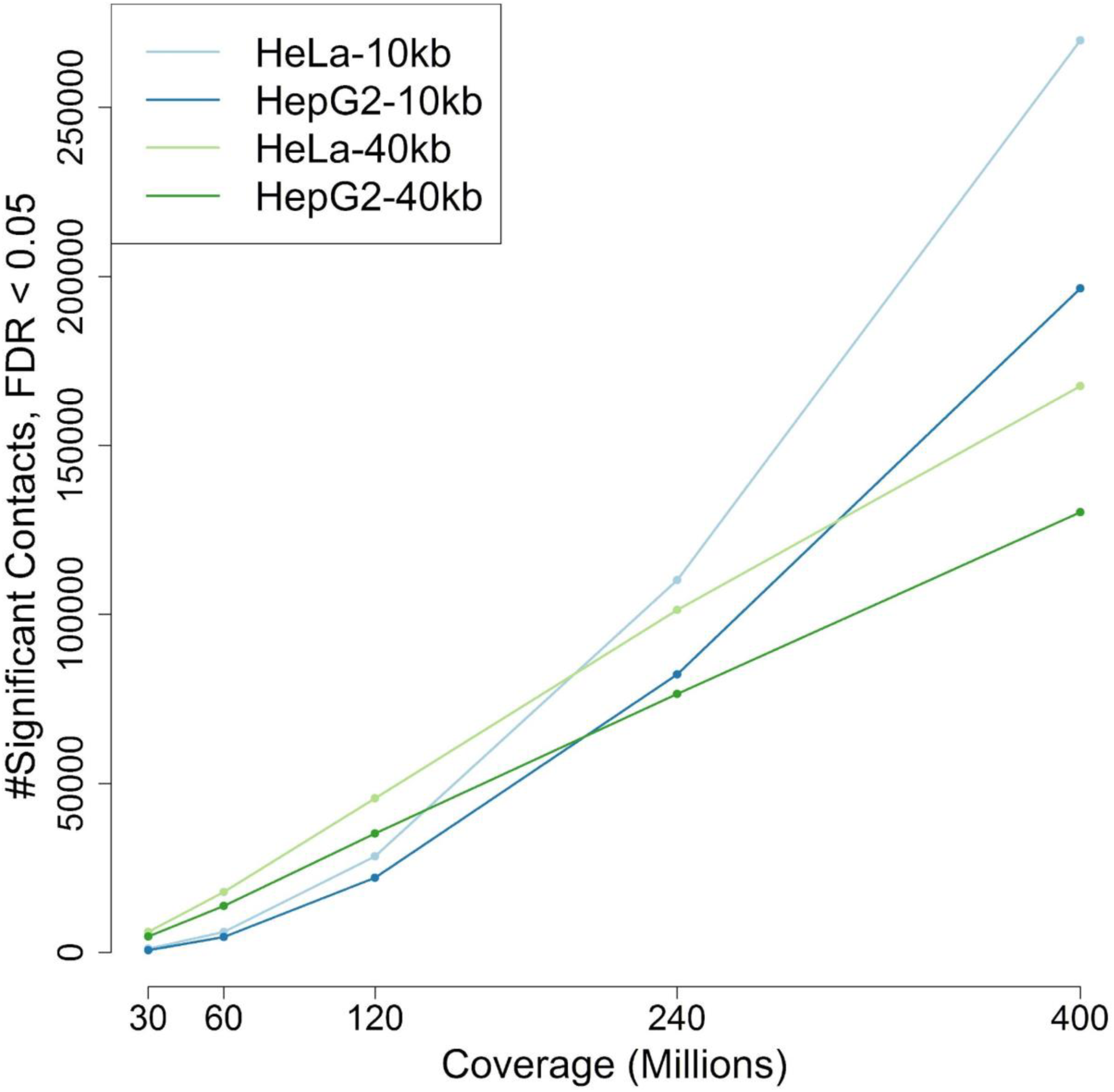
Curves showing the total number significant mid-range interactions detected by FIt-Hi-C from deeply sequenced cell types downsampled to 30, 60, 120, 240 and 400 million interactions at 10kb and 40kb resolutions.

**Supp. Fig. 16.**
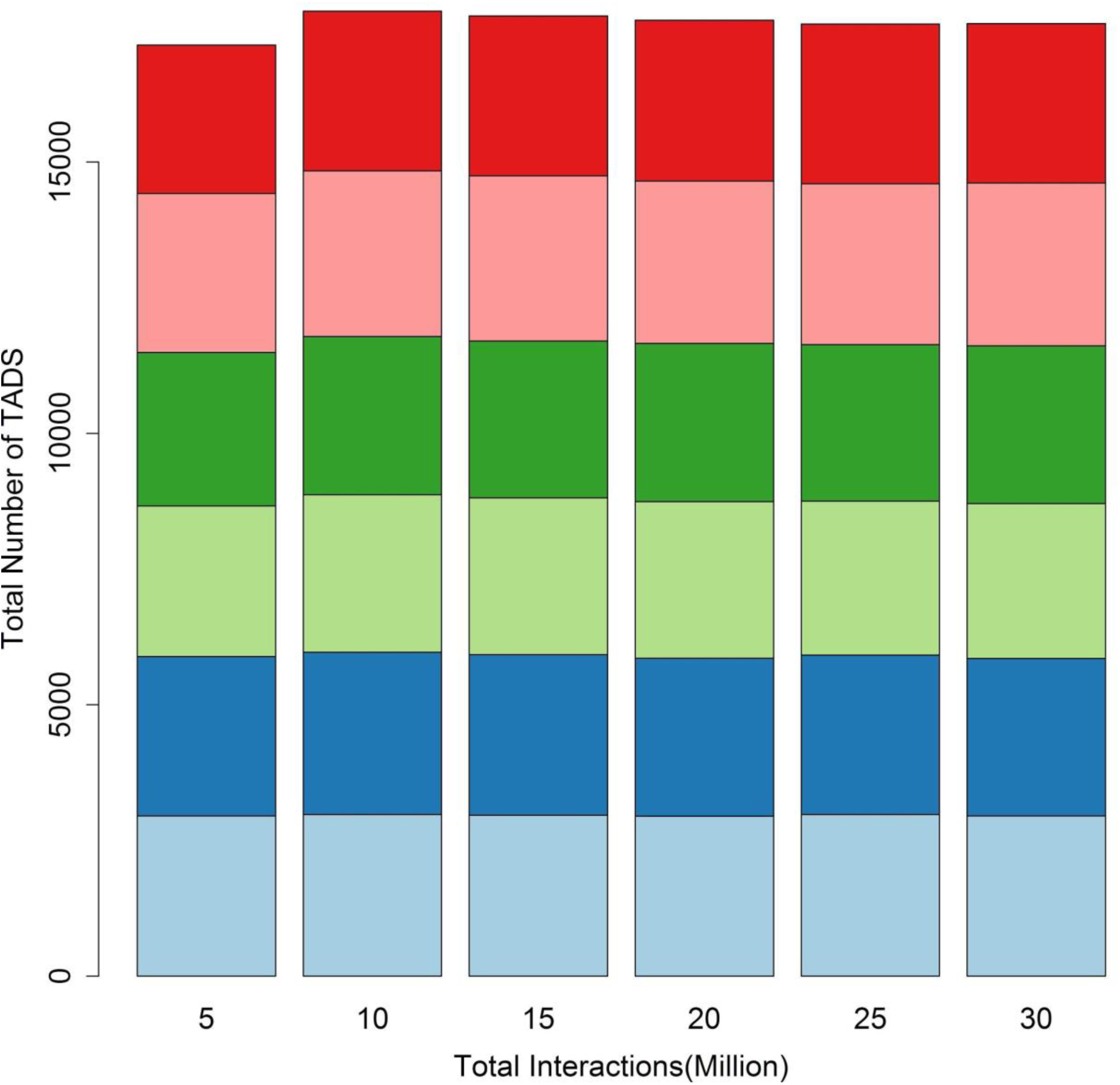
Barplots showing the number of TADs for each downsampled cell line (coded by color) at each coverage level.

**Supp. Fig. 17.**
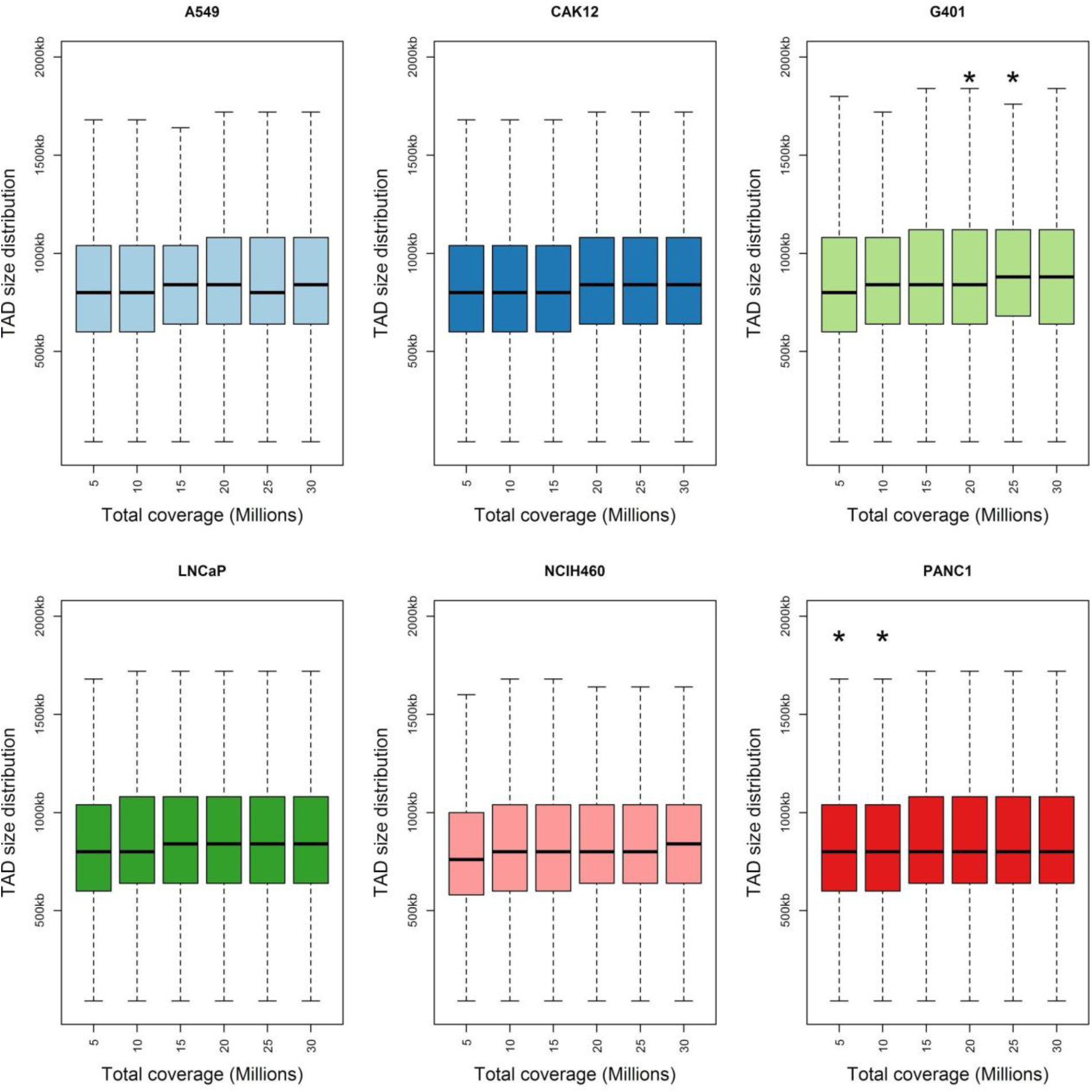
Boxplots showing the distribution of TAD sizes at downsampling level. Each plot corresponds to a simulated dataset from an individual cell type.

**Supp. Fig. 18.**
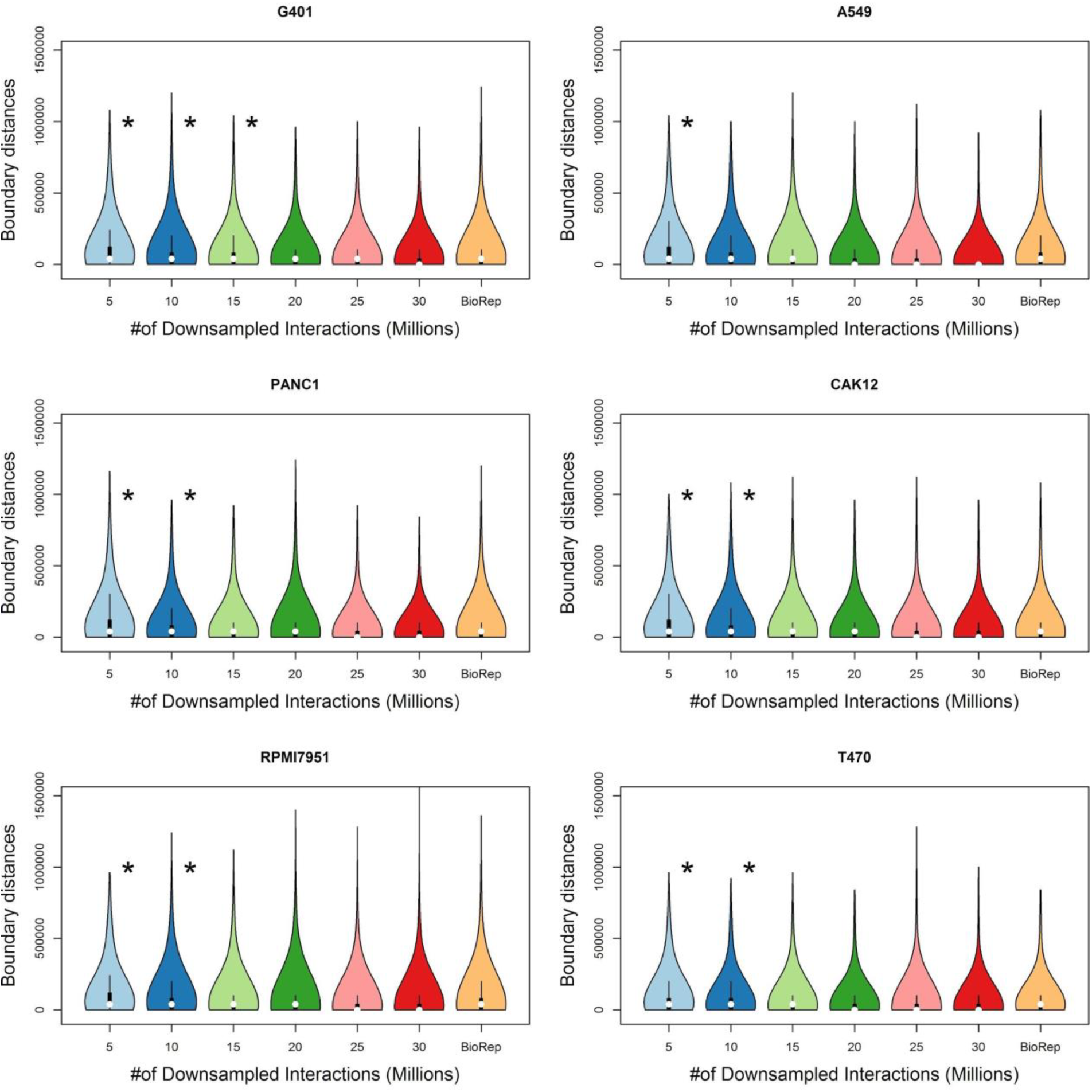
Violin plots showing the distribution of distances between domain boundaries between biological replicates and downsampled replicates. Each panel corresponds to a single cell type.

**Supp. Fig. 19.**
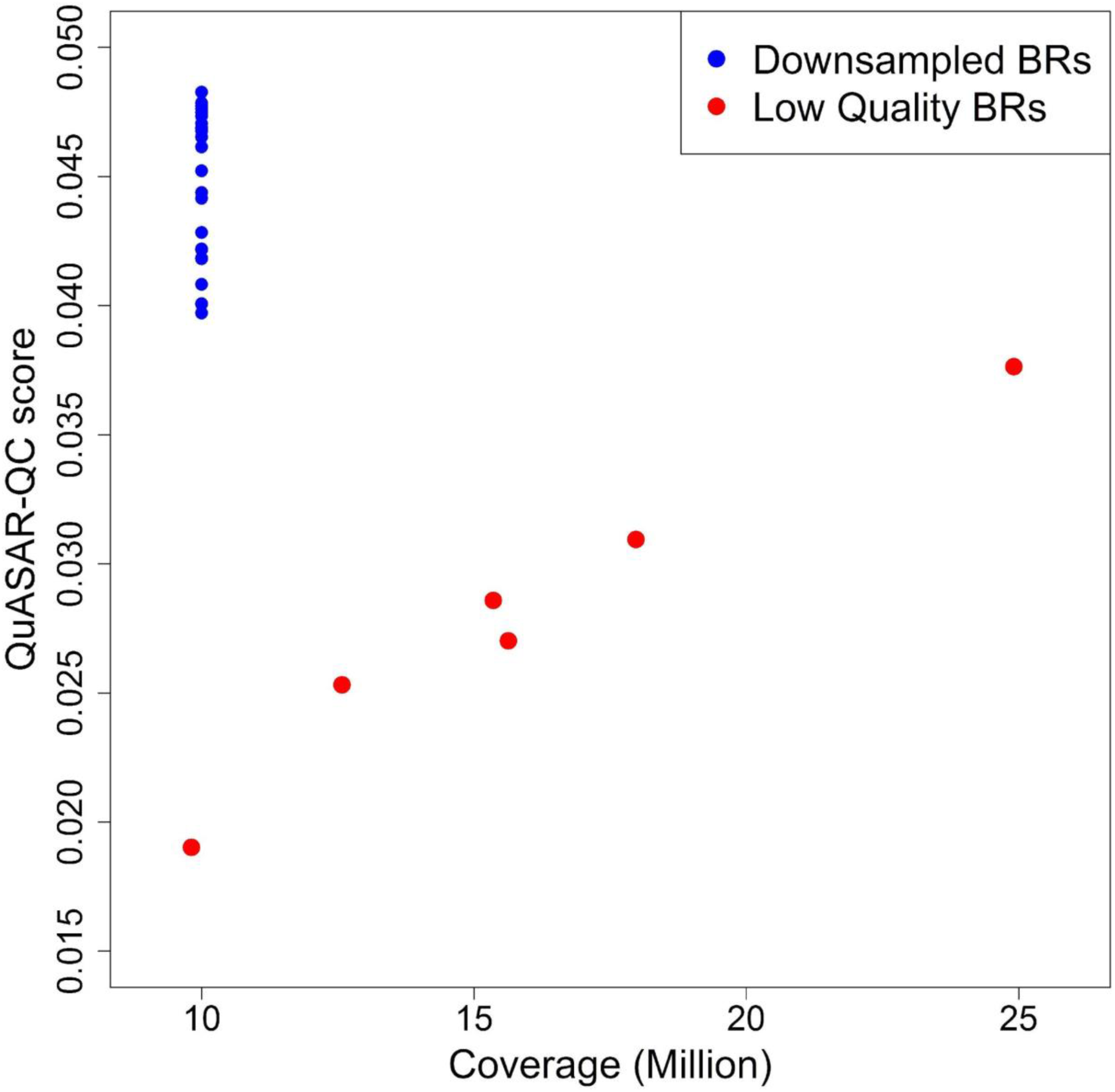
“Lower quality” data sets exhibit low QuASAR-QC scores compared to scores from downsampled data. The figure plots the QuASAR-QC score as a function of coverage. Red points are biological replicates from three cell lines designated as “low quality” Hi-C data. Blue points are from other cell types, after downsampling to 10 million interactions. Despite having higher coverage, lower quality datasets have lower QuASAR-QC scores.

**Supp. Table 1.**
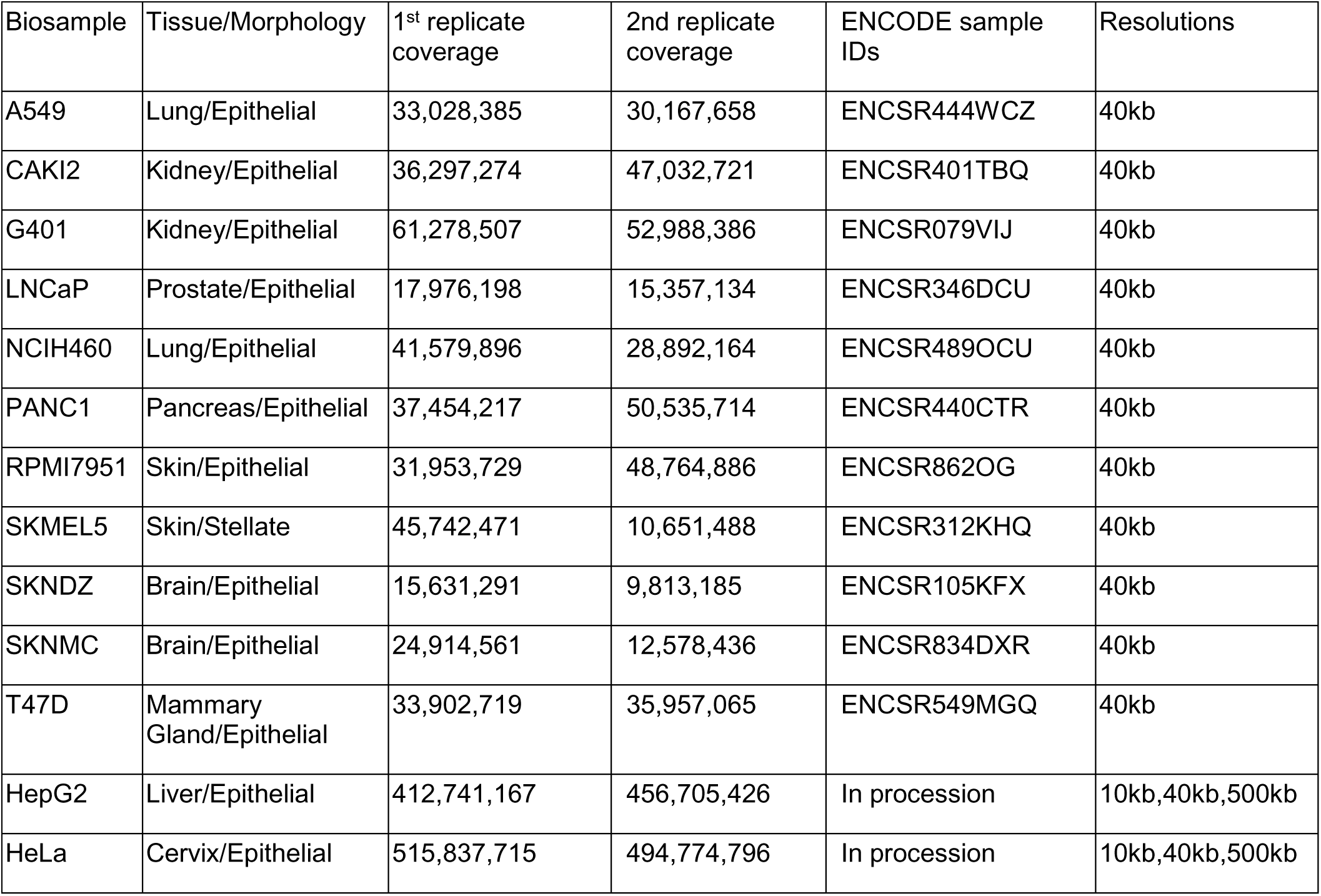
Thirteen human cancer cell types that Hi-C experiments were performed on, together with the tissue type and lineage the cells were immortalized from. Two replicate experiments were performed in each cell type. The coverage columns list the total number of intra-chromosomal interactions for the 1^st^ and the 2^nd^ replicate for each cell type. ENCODE sample ID of each experiment is provided in the corresponding column. The first 11 cell types with lower coverage vaues are binned at only 40kb resolution, whereas the last 2 cell types with large number of Hi-C interactions are binned at three different resolutions.

| Biosample | Tissue/Morphology | 1<sup>st</sup> replicate coverage | 2nd replicate coverage | ENCODE sample IDs | Resolutions |
| --- | --- | --- | --- | --- | --- |
| A549 | Lung/Epithelial | 33,028,385 | 30,167,658 | ENCSR444WCZ | 40kb |
| CAKI2 | Kidney/Epithelial | 36,297,274 | 47,032,721 | ENCSR401TBQ | 40kb |
| G401 | Kidney/Epithelial | 61,278,507 | 52,988,386 | ENCSR079VIJ | 40kb |
| LNCaP | Prostate/Epithelial | 17,976,198 | 15,357,134 | ENCSR346DCU | 40kb |
| NCIH460 | Lung/Epithelial | 41,579,896 | 28,892,164 | ENCSR489OCU | 40kb |
| PANC1 | Pancreas/Epithelial | 37,454,217 | 50,535,714 | ENCSR440CTR | 40kb |
| RPMI7951 | Skin/Epithelial | 31,953,729 | 48,764,886 | ENCSR862OG | 40kb |
| SKMEL5 | Skin/Stellate | 45,742,471 | 10,651,488 | ENCSR312KHQ | 40kb |
| SKNDZ | Brain/Epithelial | 15,631,291 | 9,813,185 | ENCSR105KFX | 40kb |
| SKNMC | Brain/Epithelial | 24,914,561 | 12,578,436 | ENCSR834DXR | 40kb |
| T47D | Mammary Gland/Epithelial | 33,902,719 | 35,957,065 | ENCSR549MGQ | 40kb |
| HepG2 | Liver/Epithelial | 412,741,167 | 456,705,426 | In procession | 10kb,40kb,500kb |
| HeLa | Cervix/Epithelial | 515,837,715 | 494,774,796 | In procession | 10kb,40kb,500kb |

**Supp. Table 2.** Empirical thresholds for distinguishing non-replicates from biological replicates for each measure at a given coverage level. Each column corresponds to empirical threshold inferred by using biological replicates and non-replicates that have been downsampled to the value in the column header (see Methods).

| | $30 \times 10^6$ | $25 \times 10^6$ | $20 \times 10^6$ | $15 \times 10^6$ | $10 \times 10^6$ | $5 \times 10^6$ |
| --- | --- | --- | --- | --- | --- | --- |
| HiC-Spector | 0.472 | 0.472 | 0.452 | 0.446 | 0.43 | 0.371 |
| GenomeDISCO | 0.833 | 0.826 | 0.815 | 0.798 | 0.767 | 0.679 |
| QuASAR-Rep | 0.63 | 0.573 | 0.497 | 0.392 | 0.253 | 0.096 |
| HiCRep | 0.882 | 0.877 | 0.868 | 0.855 | 0.829 | 0.765 |

